# Cell of origin epigenetic priming determines susceptibility to *Tet2* mutation

**DOI:** 10.1101/2023.09.04.556230

**Authors:** Giulia Schiroli, Vinay Kartha, Fabiana M. Duarte, Trine A. Kristiansen, Christina Mayerhofer, Rojesh Shrestha, Andrew Earl, Yan Hu, Tristan Tay, Catherine Rhee, Jason D. Buenrostro, David T. Scadden

## Abstract

Hematopoietic stem cell mutations can result in clonal hematopoiesis (CH) but the clinical outcomes are heterogeneous. The nature of the founder mutation and secondary mutations likely drive emergent neoplastic disease. We investigated how the state of the cell of origin where the *Tet2* mutation occurs affects susceptibility to that commonly occurring CH mutation. Here, we provide evidence that risk is written in the epigenome of the cell of origin. By characterizing cell states that underlie myeloid differentiation and linking this information to an inducible system to assess myeloid progenitor clones, we provide evidence that epigenetic markers of the cell where *Tet2* mutation occurs stratifies clonal behaviors. Specifically, Sox4 fosters a global cell state of high sensitization towards *Tet2* KO. Using GMP and primary HSC models, we show that Sox4 promotes cell dedifferentiation, alters cell metabolism and increases the in vivo clonal output of mutant cells. Our results validate the hypothesis that epigenetic features can predispose specific clones for dominance and explain why an identical mutation can result in different outcomes.

## Introduction

Upon aging, somatic cells steadily accumulate mutations at a rate of approximately 10-50 novel mutations per year across various tissues^1–6^. Every organ can be defined as a mosaic of cell clones carrying distinct mutations. Although most of these mutations are of little or no functional consequence, some may confer fitness advantage. A well-studied example can be found in the hematopoietic system, which is maintained by a pool of self-renewing multipotent hematopoietic stem cells (HSC). Upon aging, the long lifespan of HSC makes them particularly susceptible to the accumulation of genetic mutations; best estimates suggest that HSC acquire 1-3 exonic somatic mutation per cell per decade^7,8^. Among these, mutations in epigenetic regulators are commonly found in healthy elderly individuals with normal blood counts in a process called Clonal Hematopoiesis (CH), leading to an increased clonal fitness and positive selection over time^9^. CH is associated with an increased risk of all-cause mortality, a >10-fold increased risk of hematological malignancies, cardiac disease development and overall clinical adverse outcome^5,10–13^, representing a precursor state for myelodysplastic syndromes (MDS) and ultimately acute myeloid leukemia (AML)^14–16^.

One poorly understood aspect of CH is its incomplete penetrance, with most (∼96%) patients remaining disease-free over time, suggesting additional alterations to cells are needed for cell transformation. Current approaches for risk prediction are based on retrospective examination of the genetic mutations^9,17,18^. Clone size, driver gene identity, number of mutations and the fitness rate of specific variants have been demonstrated to have predictive power for transformation risk. However, while driver mutations provide useful insights into clonal progression, recent reports demonstrated that they are not complete indicators of risk. Indeed, high heterogeneity has been reported in the growth rate of clones harboring the same genetic mutation among different patients^19^ and a large proportion of CH cannot be linked to known driver genes^20^, suggesting that a large fraction of the unexplained variability can be ascribed to other cell extrinsic or intrinsic factors^21^. Environmental stimuli, such as changes in the bone marrow (BM) environment^22^, inflammation^23^ or transplant-related proliferative stress^24^ can influence clone growth. Alternatively, we hypothesize that the rate of clonal expansion can be modulated by specific properties of the clone of origin.

Using clonal tracing methods, we and others previously found that functional heterogeneity^25^ among unmutated hematopoietic clones is associated *in vivo* with defined molecular landscapes, both at the epigenetic^26^ and transcriptomic level^27^. Specifically, we demonstrated in mouse^26^ and human^28^ contexts that clonal behaviors were largely scripted in the epigenome. These clone-specific functional differences include responses to inflammatory and genotoxic stress^26^, which are preserved even upon BM transplantation in independent recipients. It is likely, therefore, that the response to a genetic mutation will similarly be influenced by the epigenome of the clone of origin.

As a model to demonstrate this concept, we investigated the epigenomic dependencies of Tet methycytosine dioxygenase 2, Tet2. Inactivating mutations to *TET2* are amongst the most prevalent in CH and are associated with the highest relative risk of disease development^14^. Mutations to this epigenetic regulator are associated with hypermethylation of cytosines at enhancer sequences and alterations in physiologic hematopoietic differentiation^29^. Knock-out (KO) mouse models accurately mirror human disease progression, with a fraction of mice transitioning from CH to different hematologic disorders months after the induction of the mutation^30,31^. Several studies have suggested that mutations to *Tet2* can have diverse functional roles across disease relevant cell types. In HSCs, *Tet2* mutations skew the HSC transcriptome towards a myelo-monocytic fate^32^ and expand the hematopoietic progenitor cell population in a cell-intrinsic manner^30^, resulting in altered differentiation and function particularly of myeloid populations^33^.

In this study, we characterize the epigenetic and functional consequences of *Tet2* inactivating mutation in early, committed and differentiated primary mouse BM populations. By pairing this information with an *ex vivo* clonal system based on Granulocytic-Monocytic Progenitors (GMP), we uncover a clone-specific epigenetic state which affects the extent of functional changes seen in myeloid cells. Further, by using HSC transplantation models we highlight that this “sensitizing” epigenetic state coincides with *in vivo* outcomes in mice, increasing the molecular and functional dysregulation associated with *Tet2* mutation. Collectively, these results provide experimental evidence that epigenetic features of the cell of origin influences the phenotype of *Tet2* mutant clonal hematopoiesis.

## Results

### Characterization of the *Cis*-regulatory landscape of GMPs *in vivo*

To study how inactivation of an epigenetic regulator may promote disease-associated programs, we performed molecular profiling at the single cell level of bone marrow (BM) hematopoietic progenitor cells from inducible *Tet2* mutant mice (Mx1-Cre Tet2fl/fl^30^) or matched controls (Tet2fl/fl) 6 months after pI:pC treatment (**Fig.1A**). Efficient Tet2 deletion was confirmed at the genomic level by ddPCR in >95% of circulating cells. Our dataset (n=4 *Tet2* KO mice, n=2 WT mice) included populations spanning from HSC to mature myeloid effector cells (Lin-progenitors, CD11b+ cells), and sorted GMPs (Lin-cKit+ Sca1-CD34+ CD16/32+), thus encompassing all the different intermediates of myeloid differentiation. GMPs represent a heterogeneous mixture of myeloid-restricted cells which connect primitive progenitors to committed multi-lineage myeloid fates^34,35^. This latter cell type is particularly relevant during oncogenic transformation where MDS/AML cells progressively co-opt GMP identities^36,37^.

**Figure 1.**
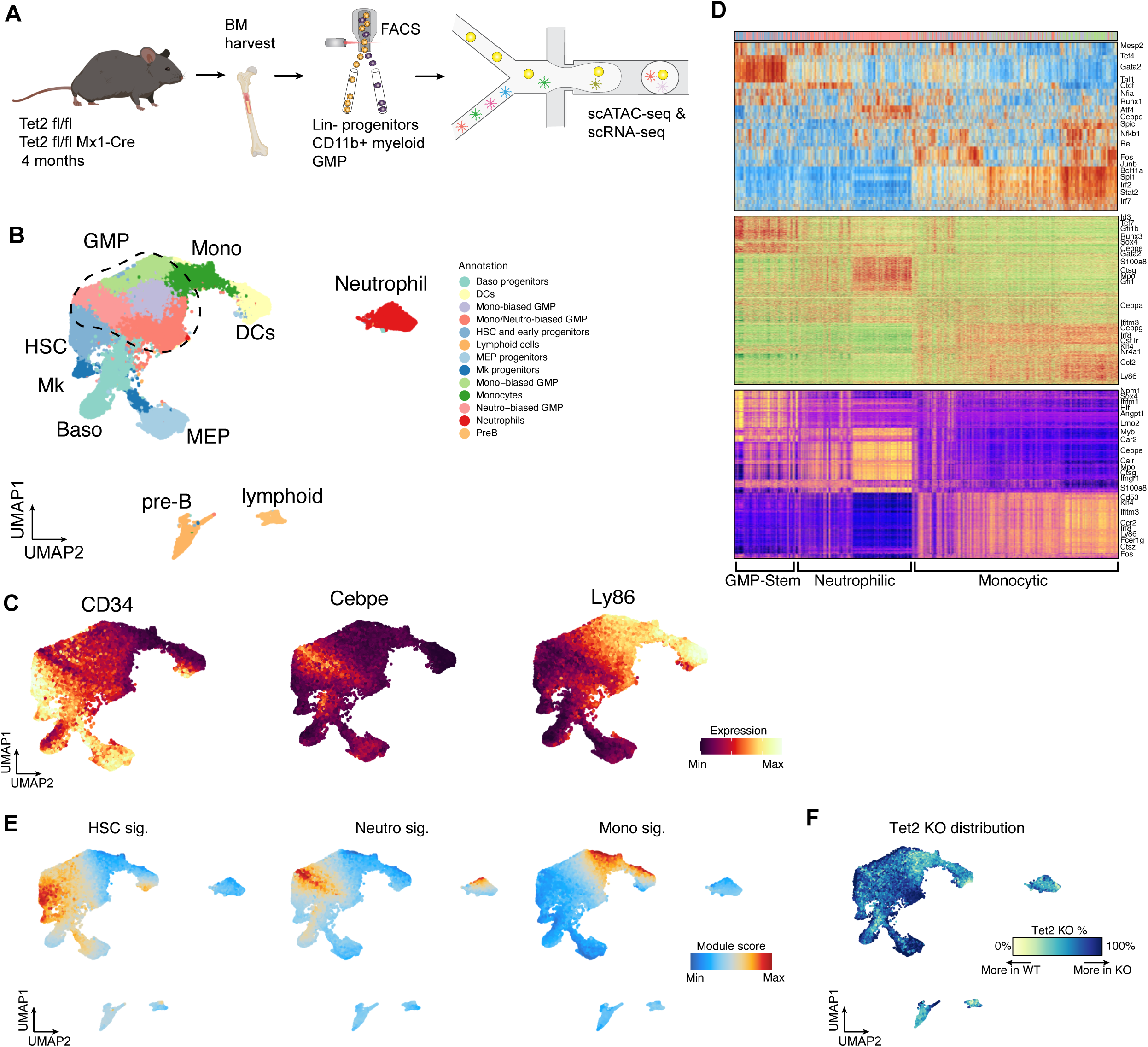
Definition of chromatin states that describe in vivo GMP differentiation. **A**) Cartoon depicting experimental strategy and molecular analyses for primary *Tet2* KO and WT populations. **B**) UMAP plot showing the combined scATACseq coordinates and annotation of sorted Lin-progenitors, CD11b myeloid cells and GMPs. **C**) UMAP plots showing RNA expression of representative markers for HSC and early progenitors (left), granulocytic (middle) or monocytic (right) differentiation. **D**) Heatmaps comparing TF accessibility(top), DORC accessibility (middle) and RNA expression (bottom) of most variable elements across GMP cells. Cells are color coded by cluster identity. **E**) UMAP plots showing cumulative accessibility for chromatin signature specific for the indicated lineages. **F**) Plot showing relative distribution of WT and *Tet2* KO cells across the differentiation trajectory.

We analyzed a total of 57,232 cells for scATAC-seq^38^ and 8,969 cells for scRNAseq and applied our previously developed analytical framework to link chromatin accessibility and gene expression across cells^39^. In this way, we were able to annotate the dscATAC-seq data using known markers for hematopoietic BM populations^34,35^ extracted from the paired single cell RNA-seq dataset (**Fig.1B, Fig.S1A-B**). Expression of representative genes for primitive or committed populations is overlaid on a 2D UMAP projection of single cells (**Fig.1C**).

To characterize the epigenetic network that regulates early progenitor cell state transitions and how this can be affected by deficiency of Tet2, we first focused on characterizing the different chromatin states that underlie GMP identity. Previous RNA-seq studies have established a hierarchical model for GMP differentiation. GMP myeloid commitment was indeed observed to start from early progenitors which co-express stem and myeloid-related genes, and then markedly bifurcate in monocytic or granulocytic fates^40,41^. We thus adopted an unbiased approach and clustered GMP based on accessibility profile for the 50 most variable transcription factor (TF) motifs after removing contaminant populations from the sorting procedure (**Fig.1D, top panel**). This analysis revealed an unexpected high degree of complexity within GMPs, suggesting the presence of multiple chromatin states. Accessibility of some key TF reportedly correlated with distinct myeloid cell identities (for example, Gata, Tal1 motifs for stem-like cells, Cebpe for granulocytic or Irf motifs for monocytic differentiation). Interestingly, high accessibility for TFs like Tcf3/4 and Runx1/2 identified states of transition between primitive GMP and more committed states, consistent with a role of these factors in regulating differentiation^35,42^. Other TFs like Nfkb, Jun, Fos, Nfia/x showed heterogeneous distribution patterns across all the different myeloid states and likely reflect activation of other regulatory properties such as inflammation and cell cycle.

In order to better identify which chromatin features can be used to identify myeloid functional identities, we took two complementary approaches. The first focused on the identification of domains of regulatory chromatin (DORCs), which prime cell identity for activation of gene expression. These regions are defined by first ranking genes based on their total number of significantly correlated peaks and focusing on genes with ≥3 associated accessibility peaks (n=3,474, **Fig.S1C,D**). Our previous work in mouse skin highlighted that DORCs are enriched in enhancers and play a key role in cell priming during lineage commitment^39^. Remarkably, DORC-associated genes identified here for hematopoietic progenitors included many known mediators of lineage specification and differentiation^34^ such as Gata1 and Mpo. Direct comparison between differential DORCs and cognate RNA expression revealed a general correlation across cells (**Fig.1D, middle and bottom panel**), and highlighted the clear presence of 3 defined GMP states (stem-like, granulocytic and monocytic). These data suggest that enhancer-like chromatin features are able to predict differentiation trajectories and cell identity during hematopoietic differentiation. Interestingly, for subsets of DORCs, we observed a gain in accessibility prior to the onset of their associated gene’s expression, consistent with a role of chromatin in priming gene expression activation^39^. For example, for Klf4, a key TF determinant of the monocytic lineage, we detected DORC activation prior to gene expression and before lineage commitment; these are quantified as ‘residuals’’ (difference of chromatin accessibility and expression of the gene^39^) (**Fig.S1E**).

The second approach is to define the core regulatory epigenetic program of the cells by aggregating accessible chromatin regions co-regulated by related sets of TFs^43^. In this manner, we identified 3 distinct signatures (co-accessibility modules, see methods) that represent relevant myeloid functional states, as shown by representing module accessibility on UMAP (**Fig.1E**) or by looking at the TF motifs and gene scores correlated to each peak module (**Fig.S1F**). An HSC signature was indeed associated with stem-like identities, as shown by gain in Runx1/2, Gata and Tal motif accessibility, whereas neutrophilic-primed signatures included gain of accessibility in the Cebpe motif and genes implicated in neutrophil differentiation (Mpo, Ctsg, Elane). The signature for monocytic priming was associated with enrichment in Irf, Klf motifs and Spi1, Ccl2, Ly86 gene scores. Accordingly, pathway enrichment analysis using top correlated genes to each signature confirmed significant association with expected functional myeloid identities (**Fig. S1G**).

Having described the presence of heterogeneous chromatin states that underline myeloid differentiation, we investigated the impact of *Tet2* mutation on the cell distribution along the trajectory. Upon *Tet2* deletion, phenotypic analysis highlighted expansion of myeloid cells in the periphery, splenomegaly, and accumulation of primitive HSC and myeloid progenitors in the BM, consistent with previous reports for this model (**Fig.S2 A-C**)^30^. In accordance with results from previous RNA-seq studies^32^, we observed that *Tet2* knockout does not result in an independent cluster of GMPs, but rather results in the redistribution of cells among the myeloid differentiation trajectory. This suggests that the presence of the Tet2 mutation skews lineage trajectories of primitive progenitors, but effects on cell identity are subtle (**Fig.1F**).

Taken together, we find GMP subsets that are epigenomically and transcriptomically distinct. We also find that *Tet2* knockout is functioning to redistribute cells among a continuum of cell states, motivating the further characterization of cell heterogeneity and their state-specific outcomes in response to *Tet2* mutations.

### *Tet2* is required for correct priming and function of myeloid progenitors

To investigate how Tet2 deficiency influences myeloid cell identity at a molecular level, we analyzed differential representations within the scATAC seq and scRNA seq datasets. Differential chromatin accessibility analysis demonstrated 2,485 peaks upregulated and 2,621 peaks significantly downregulated upon *Tet2* KO (FDR ≤ 0.01) (**Fig.2A).** To confirm that these chromatin changes are related to direct binding of Tet2, we overlapped the differential peaks with available *Tet2* ChIP-seq data from GMP-relevant hematopoietic cells^44^. We indeed found overlap for a high fraction of differential accessibility peaks (Fisher exact-test *P* < 0.001), finding 30.9% for peaks repressed and 21.8% for peaks induced upon Tet2 KO. In accordance with the *Tet2* function as a DNA demethylase^29^, a larger fraction of accessible regions predicted to be bound by Tet2 are observed to be repressed upon *Tet2* KO. In order to evaluate the level of epigenetic dysregulation on a single cell basis, we calculated single cell scores for accessibility of *Tet2* differential peaks from Fig.2A (**Fig.2B**). Intriguingly, we observed a high spread in the distribution of *Tet2* KO cells as compared to WT ones, suggesting that there is a level of heterogeneity in how single GMP cells are epigenetically affected by Tet2 mutation.

**Figure 2.**
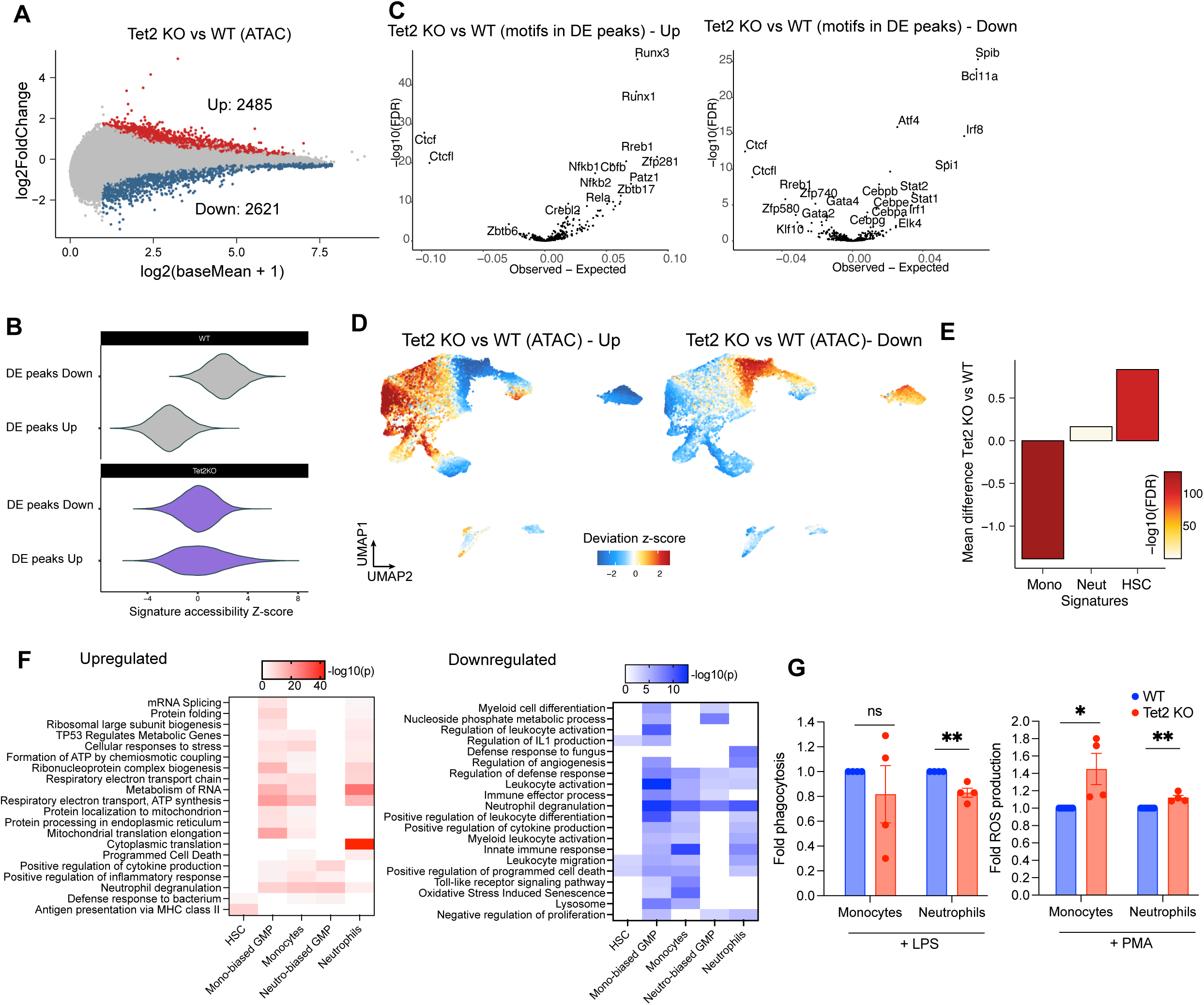
*Tet2* is required for correct myeloid cell priming. **A)** Plot showing the fold change in the accessibility of chromatin peaks comparing WT and *Tet2* KO GMP cells. Significantly upregulated peaks are colored in red while significantly downregulated peaks are colored in blue. **B**) Z-score accessibility of DE peaks from Fig. 2A measured per-cell relative to the entire dataset. Genotype is indicated. **C**) TF motif accessibility comparing WT and *Tet2* KO GMP cells measured in differential peaks regions (Left: upregulated,, Right: downregulated). **D**) Cumulative accessibility of differential upregulated (left) and downregulated (right) peaks plotted on UMAP coordinates comparing WT and *Tet2* KO GMP cells. **E**) Plot showing the mean difference in chromatin signatures comparing WT and *Tet2* KO GMP cells. Level of significance is color coded. **F**) Pathway enrichment analysis of upregulated (left) and downregulated (right) genes in the indicated cell populations. Significance is color coded. **G**) Fold change in the level of phagocytosis (left) or production of ROS (right) comparing WT and *Tet2* KO primary monocytes and neutrophils stimulated as indicated (n=4, p<0.01).

We then further investigated the functional relevance of the chromatin regions differentially regulated after *Tet2* knockout. To do this, we calculated enrichment of TF motifs among differentially accessible chromatin peaks and used these as an indicator of potential TF activity. We detected significant enrichment of many TF families within differentially regulated peaks comparing WT and *Tet2* KO GMP cells (permutation *Z*-test FDR < 0.001; **Fig.2C**). Specifically, among the activated peaks, we observed a strong enrichment of Runx1/2 motifs, likely indicating an effect of *Tet2* on cell differentiation at early progenitor stages. The repressed peaks, in contrast, were enriched for Bcl11a, Irf8, Stat, Atf4 motifs, TFs that regulate monocytes and granulocytes cell commitment and differentiation. Interestingly, across both upregulated and downregulated peaks inflammation related TF motifs (Nfkb for activated, Irf1/8 for repressed) were identified, consistent with an enhanced inflammatory state as previously observed in *Tet2* KO models^45,46^. These alterations collectively highlight an impairment of myeloid differentiation in response to *Tet2* knockout. We also consistently observed a decrease in Ctcf motifs, which are sensitive to DNA methylation. This finding correlates with reduced CTCF binding previously observed in IDH mutant gliomas^47,48^, and might indicate overall disruption of chromatin insulation in *Tet2* KO in the hematopoietic context.

As a complementary approach, we assessed the differentiation stage where *Tet2*- differential chromatin regions are activated. Remarkably, peaks induced upon *Tet2* KO showed high accessibility in primitive cells, suggesting that Tet2 participates in chromatin regulation to affect HSC and progenitor commitment. Peaks repressed upon *Tet2* KO showed highest accessibility upon progenitor commitment towards myeloid fates (**Fig.2D**). These data are consistent with *Tet2* being required for priming of GMPs towards both monocytic and granulocytic lineages. To validate these results in an orthogonal way, we analyzed our previously defined signatures for HSC and myeloid cells comparing WT and *Tet2* KO cells (**Fig.2E**). Upon Tet2 KO, we observed significant loss in accessibility for the monocytic signature and an increase in accessibility in the HSC signature, further indicating that chromatin changes mediated by Tet2 alter the core regulatory network at different levels of myeloid differentiation.

Since our differential peaks analysis indicates extensive epigenetic remodeling, we analyzed the downstream alterations at the level of gene expression. Enrichment analysis was conducted for cell subsets covering the key stages of myeloid differentiation (**Fig. 2F**). Among the upregulated pathways, gene sets were identified that broadly associate with altered oxidative state, translation, response to cell pathogens and inflammation across the granulocytic and monocytic cell fates. Intriguingly, among the most significant terms for HSC we found MHC class II antigen presentation, in accordance with this recently described immunosurveillance mechanism being used in safeguarding pre-malignant states^49^. Downregulated terms indicate a differentiation arrest of GMP as well as substantial alterations in physiologic myeloid activation and functions in the absence of *Tet2*.

Finally, to validate whether the priming defect we observed in GMP cells upon *Tet2* KO leads to functional alterations after differentiation, we analyzed cell functionality of monocytes and neutrophils with inflammatory stimuli. Mature *Tet2* KO myeloid cells from both lineages showed increased reactive oxygen species (ROS) production upon treatment with PMA (**Fig.2G**). This finding is consistent with a described role for Tet2 in repressing inflammatory gene expression^45^. Interestingly, we also observed decreased phagocytosis in *Tet2* KO cells (**Fig.2G**), suggesting that impaired microbial clearance of monocytes and neutrophils might play a role in the amplification of inflammatory responses observed in *Tet2* KO. Indeed, a previous study highlighted that bacteria accumulation (resulting from a dysfunctional intestinal barrier) is critical for sustaining IL6-induced myeloproliferation observed in *Tet2* KO model^50^. Similar functional impairment has also been reported in a *Tet2* deficient zebrafish model^51^.

Overall, these data shed light on the *Tet2*-related molecular mechanisms involved in correct myeloid cell priming that influence downstream effector functions.

### Linking epigenetic states to functional properties of GMP clones

To shed light on the epigenetic heterogeneity observed in primary progenitors (Fig.2B), we thus sought to determine how diverse GMP cell states may be differentially altered in their function and in response to *Tet2* knockout. We hypothesized that individual GMP clones may stably inherit epigenetic, transcriptomic and functional states. To perform paired functional and molecular analyses on single hematopoietic clones, we took advantage of a high-throughput *in vitro* model of primary myeloid progenitors conditionally immortalized using HoxB8 retrovirus fused to Estrogen Receptor domain (Hoxb8-ER)^52^. This culture system enables extended self-renewal of primary GMPs, allowing their single-cell isolation and indefinite expansion as pure clones. Upon withdrawal of the estrogen analogue, the clones undergo physiological differentiation into mature myeloid cells with intact molecular characteristics and *in vivo* functions of primary cells^53^. We generated a dataset of 28 individual clones from a *Tet2* fl/fl mouse and each of them was divided into cohorts for induction of *Tet2* KO or control (**Fig.3A**). Each clone pair is derived from an individual GMP cell. We could confirm nearly complete introduction of *Tet2* targeted deletion after Cre transduction in all the clones using a quantitative PCR assay (mean 97%, **Fig.S3A**). 24 clone pairs were generated in the presence of SCF, which primes them towards granulocytic fate, and 4 of them were derived in the presence of GM-CSF which arrests them at a more mature stage and directs them towards a monocyte-macrophage fate (**Fig.S3B,C**). We profiled the epigenetic landscape of the clones by performing multiplexed bulk ATAC-seq. Our analyses revealed highly heterogeneous patterns of chromatin accessibility (**Fig.3B**), with separate clustering of SCF and GM-CSF clones.

**Figure 3.**
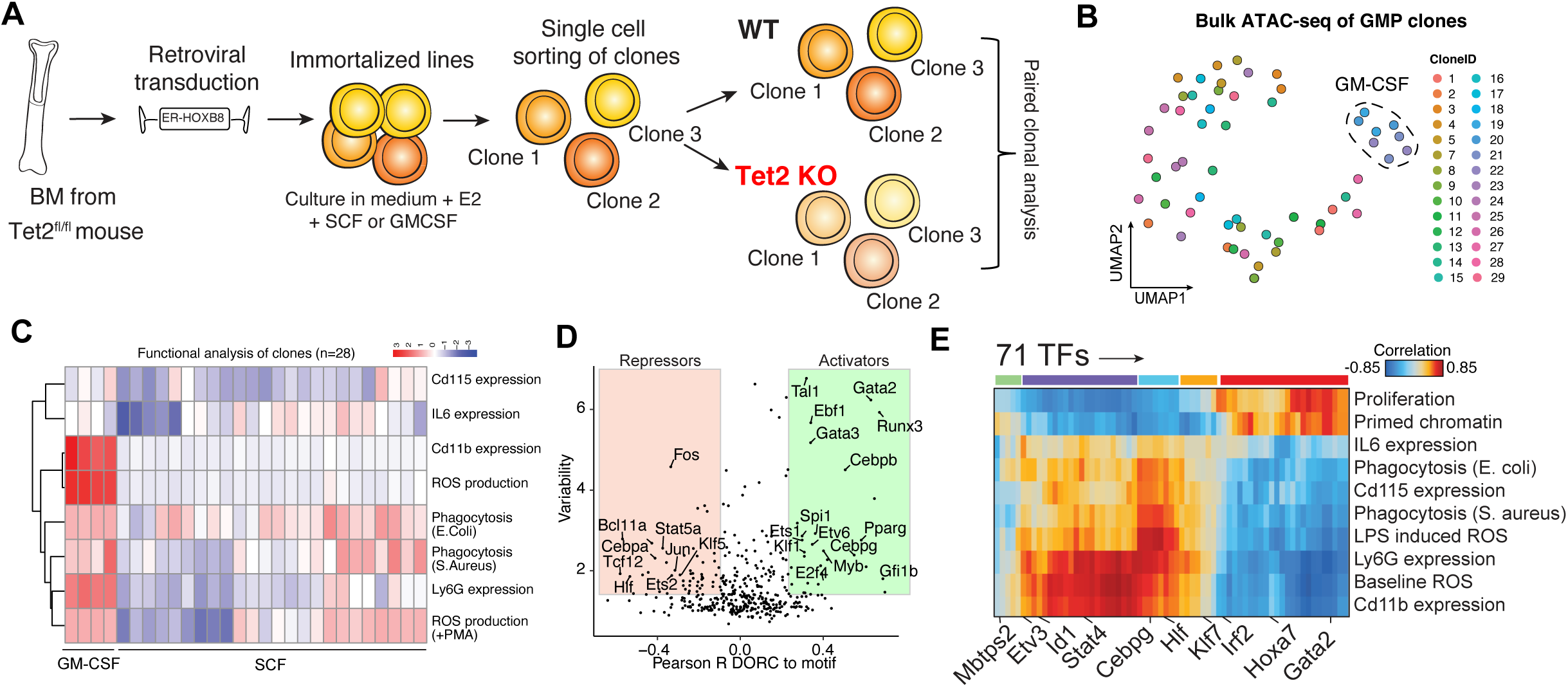
GMP system as a model for heterogeneity reveals clone-specific changes. **A)** Cartoon describing experimental workflow for the generation of paired WT and *Tet2* KO single GMP clones using HoxB8-ER system. **B**) UMAP plot showing heterogeneity in the chromatin profile of different clones by high throughput bulk analysis. Clones within the dashed line were generated using GM-CSF whereas the rest of the clones were generated with SCF. **C**) Heatmap showing performance of different WT GMP clones tested the indicated in vitro functional assays. **D**) Plot showing Pearson correlation between chromatin accessibility of DORC and respective TF motifs in GMP clones. Positive correlation (green) indicates activators whereas negative correlation (red) indicates repressors. **E**) Heatmap showing Pearson correlation between TF activity measured by motif accessibility and cell function of GMP clones measured by the indicated functional assays.

We then characterized cell function by performing multiple *in vitro* functional assays on the differentiated progeny derived from clonal progenitors (**Fig.3C, Fig.S3D-G**). We measured the expression of myeloid markers, production of inflammatory cytokines (IL6), cell competency to clear pathogens in basal and stimulated conditions by ROS generation and phagocytic ability. Unsupervised clustering revealed the presence of distinctive functional groups (**Fig.3C**), supporting HoxB8-ER as a powerful model for investigating myeloid clonal heterogeneity. Overall, GM-CSF clones showed increased potency in accordance to their higher basal differentiation state. SCF clones showed high heterogeneity of behavior, with some clones showing potent effector functions while others were substantially less proficient.

We next sought to determine whether these functional alterations may be explained by heritability of chromatin states at the progenitor level. To do this, we correlated accessibility of variable TF motifs with those of their cognate DORCs. This analysis identified activators and repressors which regulate chromatin accessibility including many known TFs involved in differentiation. These include Runx1, Hlf and Tal1; GMP specification, such as Cebpa-b; effectors of myeloid cell function, such as Nr4a1-3 and members of the Atf family^54,55^ (**Fig.3D**). Importantly, performing correlation analyses between the functional data and significant TF regulators, we could directly link the epigenetic states of the GMP progenitors to their proficiency in performing effector functions (**Fig.3E**). We observed high correlation with proliferative potential and in vivo output capacity with stem-associated transcription factors (members of the Runx, Gata, Hoxa families, Sox4 and Tal1). Most effector functions correlated with Nr4a1, Atf, Cepb and Stat family members. IL6 and CD115 expression mostly correlated with Nfkb2 and Klf5, consistent with their described role in inflammatory responses^56,57^.

Collectively, these data show that the GMP clonal system is a robust model that enables the study of myeloid heterogeneity and directly links epigenetic features with relevant cell functions. It further highlights that the differentiation fate of single clones is scripted in the chromatin accessibility profile of the cell of origin.

### Paired clonal analysis defines clone-specific sensitization to *Tet2* mutations

We then investigated whether different GMP clones have different levels of molecular perturbation after *Tet2* KO. The identification of molecular determinants of sensitivity to the mutation might inform which clones will be more affected in their function (and thus likely to drive disease development). We thus investigated functional alterations after differentiation of WT and mutated clone pairs. *Tet2* mutated clones showed major functional alterations as compared to WT sister counterparts (**Fig.S3D-G**). There was significant alteration in myeloid differentiation marker expression, reduction of phagocytosis of E.Coli bacterial particles, increased production of ROS and production of IL6 upon PMA and LPS stimuli. The alterations collectively indicate hyperresponsiveness to inflammatory stimuli and impaired host anti-bacterial defense. Importantly, these results mirror functional changes observed in primary *Tet2* KO cells *in vivo* by previous reports^46,58,59^ and in our own primary dataset of primary monocytes and neutrophils (Fig.2G), Importantly, we found that some clones had a significantly altered phenotype in response to *Tet2* KO while others were largely unperturbed in their functions. We thus summarized the extent of functional perturbation of each pair of sister WT and *Tet2* KO clones by calculating a “Functional Perturbation Score” (**Fig.4A**), computed as the euclidean distance of each clone pair including all the above mentioned functional *in vitro* data (Fig.S3D-G). GM-CSF derived clones were much less perturbed in their behavior as compared to SCF ones, possibly because these cells are immortalized at a differentiated stage whereas our in vivo data suggest *Tet2* is mostly required earlier in the hematopoietic hierarchy. Among SCF clones, some clones show a higher extent of functional alteration in the absence of *Tet2*, and overall, high perturbation correlated with lower differentiation properties (**Fig.S4A**). As an additional functional parameter of self-renewal ability, we tested the *in vivo* output of the GMP clones by transplantation in lethally irradiated recipients. SCF-derived GMP clones differentiate *in vivo* and give rise to a transient wave of granulocytic progeny in the peripheral blood (PB) in accordance with their limited self renewal potential (**Fig.S4B**). Notably, *in vivo* output of the transplanted clones was collectively higher for *Tet2* mutants and we observed again clone-specific divergent responses (**Fig.4B**). Collectively, these data demonstrate that Tet2 largely impacts *in vitro* and *in vivo* functions of GMPs, influencing preferentially the behavior of some clones and not others, often linked to the differentiation state.

**Figure 4.**
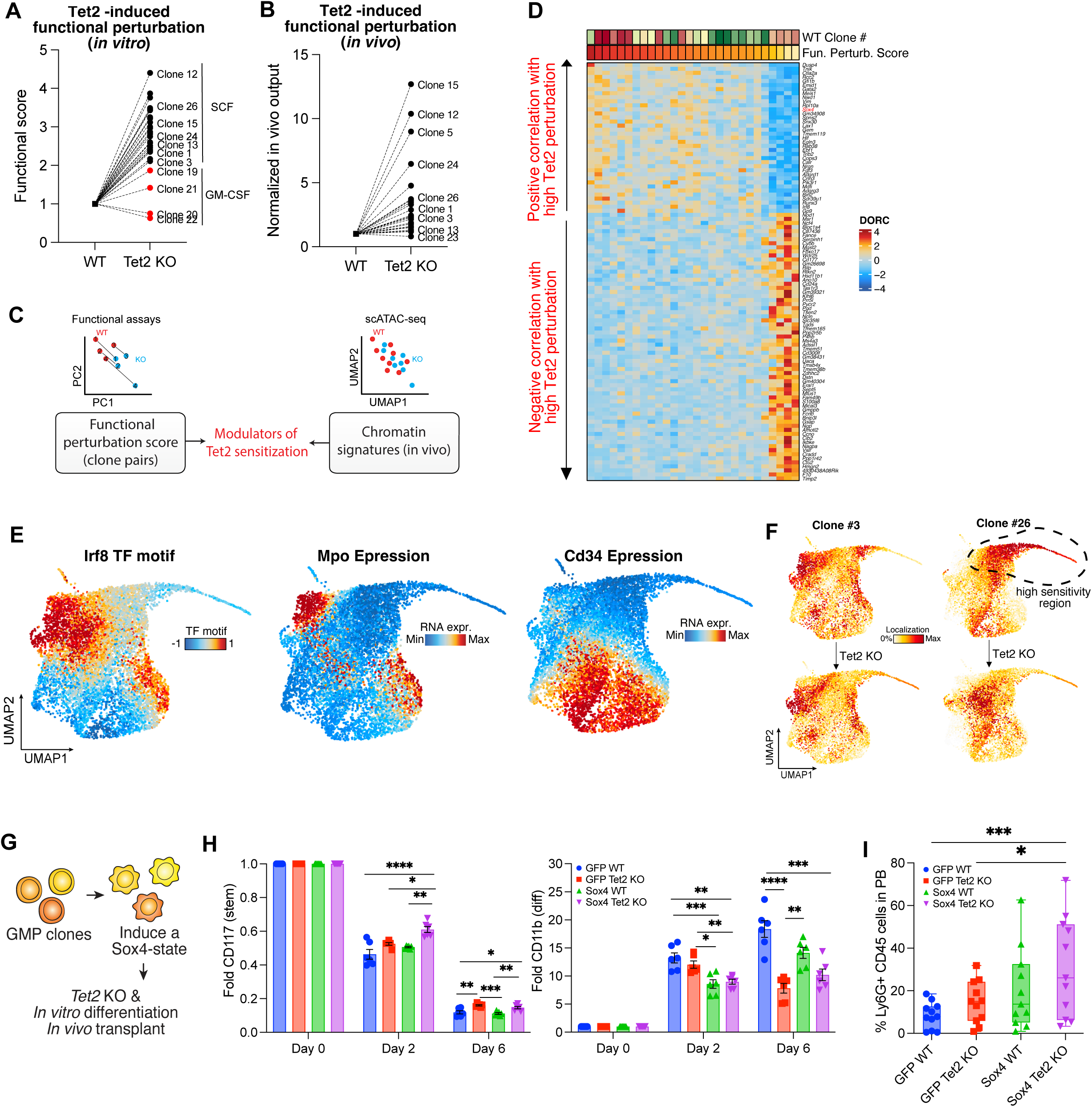
GMP clonal system defines sensitization status to *Tet2* mutation. **A**) Functional perturbation score for each WT and *Tet2* KO GMP sister clones, indicating the extent of functional alterations measured using *in vitro* assays **B**) *In vivo* output for each WT and *Tet2* KO GMP clone pair, indicating the extent of functional alterations measured upon *in vivo* transplantation. **C)** Schematics depicting the analysis framework utilized to link clone phenotypes to *in vivo* chromatin states and identity epigenetic predictors of *Tet2* sensitization states. **D**) Heatmap showing the top DORC chromatin regions correlated to GMP clones functional score. **E)** UMAP plots showing representative markers (RNA or ATAC) relevant for GMP identity and differentiation. **F**) UMAP plots showing single cell distribution of WT and mutated GMP clone pairs analyzed by SHARE seq. **G**) Schematics representing the experimental workflow and analysis for generating Sox4-overexpressing GMP clones. **H**) Expression levels of Kit (left) and Cd11b (right) at different time points during GMP clone differentiation. Values are normalized to day 0. n=6, p<0.05, p<0.01, p<0.001, p<0.0001. **I**) *In vivo* output of GMP clones treated as indicated. PB analysis is performed at day 7 after GMP transplantation. n=12, p<0.05, p<0.001

Since we have shown that the GMP system is suitable to directly correlate epigenetic state and cell function, and that different clones are functionally impacted by *Tet2* KO to different extents, we investigated whether there are epigenetic states which affect the *Tet2*-related response. We set up an analytical framework to predict sensitivity to *Tet2* KO by correlating functional responses in paired KO and WT clones and differential signatures defined in our *in vivo* primary dataset (**Fig.4C**). We ranked WT clones by Functional Perturbation Score, and assessed correlations of accessibility of chromatin loci with regulatory activity (DORCs) in primary cells. This genome-wide unbiased analysis defined DORCs significantly enriched in sensitized clones (**Fig.4D**), and among them were TFs regulating stem cell identity, such as Gata2, Meis1, Gfi1b and Sox4. These data suggest that a stem-like state in GMP clones might sensitize them to *Tet2* mutation.

We next sought to better characterize whether all cells in the clone were sensitized to *Tet2* KO. We thus performed combined scRNAseq and ATACseq analysis using SHARE-seq^39^ on a subset of undifferentiated GMP clones characterized by different properties. Visualization of some key motifs and marker genes allowed us to identify regions characterized by different levels of activity of key myeloid differentiation regulators (**Fig.4E**). By analyzing the distributions of single cells on the UMAP coordinates, we observed differential localization across different WT clones (**Fig.S4C**). This result validates that functional heterogeneity observed upon differentiation (Fig.3C) is preceded by extensive molecular priming at the level of progenitor cells. Intriguingly, by performing a paired analysis and considering the SHARE-seq profiles of the *Tet2* mutated clones, we observed a marked perturbation in cell state upon *Tet2* KO (**Fig.4F, Fig.S4C**). Clones showing high Functional Perturbation Score, such as 12 and 26, were enriched in a separate region and showed a marked shift in UMAP coordinates after *Tet2* KO as compared to other analyzed clones. These data reinforce the notion that a permissive epigenetic state drives higher molecular perturbation upon *Tet2* KO.

We next tested whether the activity of chromatin regulators positively correlated with high *Tet2* sensitivity might be functionally implicated to foster the permissive clonal state. Among the positively correlated TFs from Fig.4D, *Sox4* emerged of particular interest given its known functional association with AML when overexpressed with multiple oncogenes^60–63^. Sox4 is a member of the SRY-related HMG-box TF family. Its expression is highest in primitive progenitors and progressively decreases with myeloid differentiation^61^. Its upregulation has been associated with increased HSC activity and self-renewal^64,65^ and blocking differentiation during leukemogenic transformation^66^. ATAC-seq tracks at the proximal promoter region of Sox4 confirmed higher accessibility in clones showing a high sensitization profile (**Fig.S4D).**

Here, to match physiologically relevant levels of Sox4, we overexpressed its ORF using a lentiviral construct with mild promoter activity in hematopoietic cells^67^(**Fig.4G**). With this system, we achieved a 2-fold increase in *Sox4* expression in GMP clones as compared to endogenous expression levels (**Fig.S4E**). We measured the functional effects of Sox4 overexpression including a time course of GMP differentiation using surface expression of c-Kit and Cd11b. While *Tet2* KO resulted in a decrease of CD11b and an increase in Kit levels compared to control, concomitant overexpression of Sox4 delayed GMP differentiation especially in the early time points (**Fig.4H**). This differentiation delay is not due to reduced cell proliferation, but likely to increased self-renewal divisions, as Sox4 overexpression increased proliferation of WT or *Tet2* KO GMP clones both in steady state and upon differentiation (**Fig.S4F-G**). Finally, we measured GMP clonal output by mouse transplantation assays. Notably, the combination of Sox4 overexpression significantly increased the myeloid output of *Tet2* KO GMP clones *in vivo* (**Fig.4I**).

Collectively, these data demonstrate that defined epigenetic states of clones can lead to divergent phenotypes in response to an identical genetic perturbation. Moreover, Sox4 activity promotes a sensitized cell state to *Tet2* which is characterized by stem-like properties.

### *Sox4* enhances epigenetic dysregulation observed after *Tet2* deletion

The GMP clones are useful to study clone-specific responses and the functional impact of *Tet2* on myeloid lineage progenitors, the target cell population of CH is more primitive. We therefore further validated our findings in a primary HSC system (**Fig.5A**).

**Figure 5.**
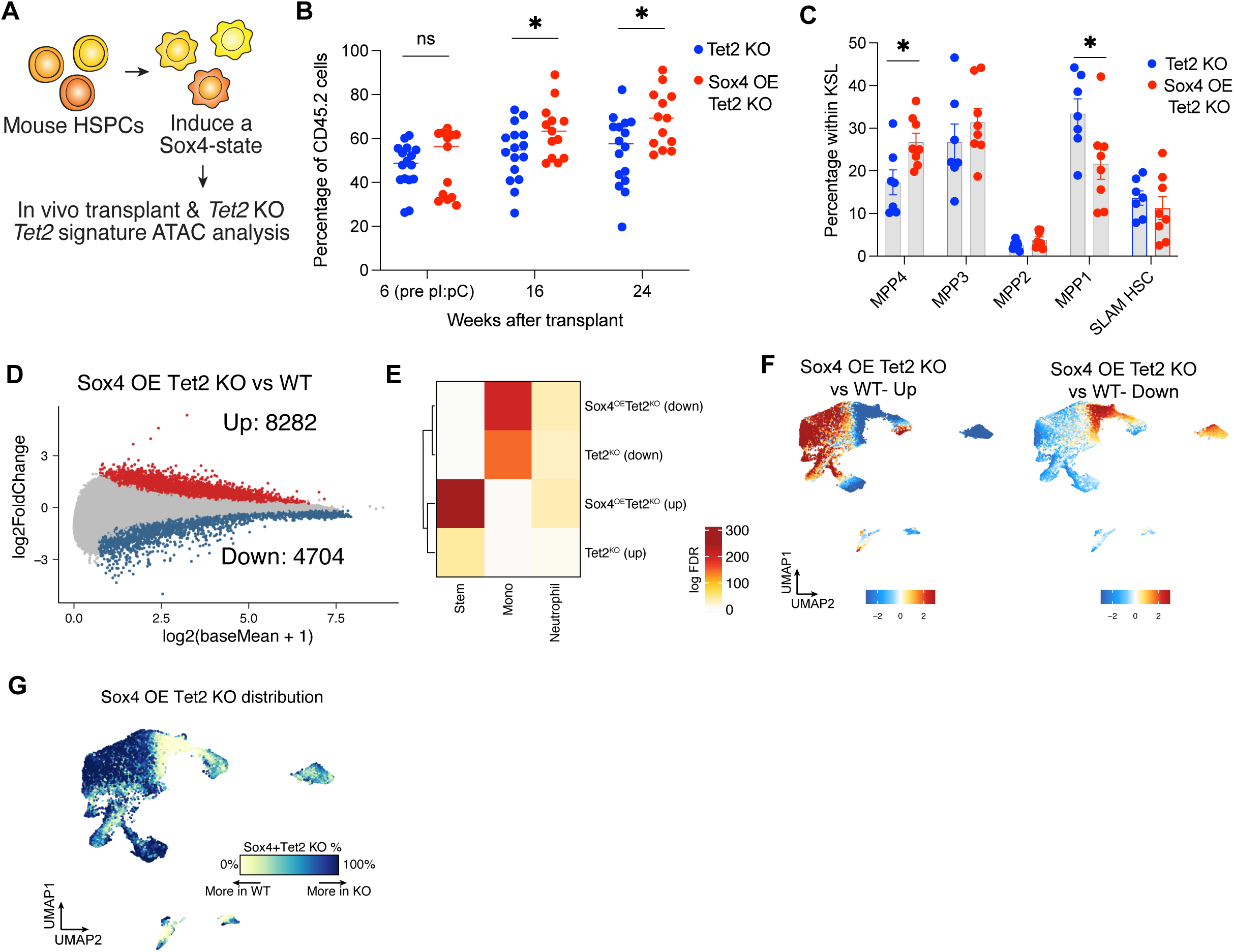
*Sox4* enhances epigenetic dysregulation observed after *Tet2* KO in primary cells. **A)** Schematics representing the experimental workflow and analysis for generating Sox4-overexpressing primary HSPC in *Tet2* fl/fl background. **B**) Percentage of CD45.2 cells measured at different time points in the PB of mice transplanted with inducible *Tet2* KO cells together with overexpression of GFP (Tet2 KO, n=16) or Sox4 (Sox4 OE *Tet2* KO, n=13). p<0.05. **C**) Distribution of the indicated populations within Lin-cells in the BM of mice from B 35 weeks after transplant. Populations are defined as in^69^. **D**) Plot showing the fold change in the accessibility of chromatin peaks comparing WT and Sox4 OE *Tet2* KO GMP cells analyzed by scATACseq. Significantly upregulated peaks are colored in red while significantly downregulated peaks are colored in blue. **E**) Plot showing the overlap between chromatin signatures and differential peaks comparing *Tet2* KO and Sox4 OE *Tet2* KO GMP cells. Level of significance is color coded. **F**) Cumulative accessibility of differential upregulated (left) and downregulate (right) peaks plotted on UMAP coordinates comparing WT and Sox4 OE *Tet2* KO GMP cells. **G**) Plot showing relative distribution of WT and Sox4 OE *Tet2* KO cells across the differentiation trajectory.

Primary KSL (Kit+ Sca-1+ Lin-) cells from *Tet2* fl/fl Mx-1 Cre mice were transduced with either GFP or Sox4 overexpressing lentiviral constructs and transplanted into lethally irradiated recipients in competition with unmanipulated CD45.1(STEM)^68^ congenic cells. *Tet2* deletion was introduced 6 weeks after the transplant by pI:pC administration and the *in vivo* contribution was measured up to 24 weeks post transplant. We observed a significant increase in the *in vivo* repopulation of Sox4 OE *Tet2* KO cells as compared with *Tet2* KO alone (**Fig.5B**). Mice transplanted with Sox4 OE HSC did not show a repopulation advantage as compared to controls, validating that Sox4 specifically provides HSC competitive fitness only when in combination with *Tet2* deletion (**Fig.S5A**). We then investigated if Sox4 overexpression induced changes in the cell composition and validated that Sox4 OE *Tet2* KO mice did not show any significant skewing in the distribution of the differentiated BM lineages (**Fig.S5B**) and the Lin-progenitors (**Fig.S5C**). Significant differences were instead observed in the distribution of KSL primitive cells, with an expansion of MMP4 cells (**Fig.5C**). This population represents a highly proliferative direct precursor of both lymphoid and myeloid progenitors^69^ and its accumulation is consistent with an exacerbation of impaired myeloid priming observed in the *Tet2* KO context.

In order to characterize the molecular changes associated with the interaction between *Sox4* and *Tet2*, we performed dscATAC-seq on BM populations with the same sorting strategy as in Figure 1 (HSC to mature myeloid effector cells, n=2 *Tet2* KO, n=2 Sox4 OE *Tet2* KO). We performed differential chromatin accessibility on the GMP subset and found 8282 peaks upregulated and 4704 peaks significantly downregulated in the Sox4 OE *Tet2* KO condition relative to WT cells (**Fig.5D**). Overlapping the differential peak signatures, Sox4 OE *Tet2* KO included the vast majority of differential peaks observed in the Tet2 KO alone samples (82% of downregulated and 91% of upregulated peaks) (**Fig.S5D**). In addition, 6019 peaks were uniquely upregulated comparing Sox4 OE *Tet2* KO condition relative to WT cells, consistent with an established role for Sox4 as a transcriptional activator^70^. Overlapping the differential peaks with the previously identified epigenetic signatures that define myeloid differentiation, demonstrated that Sox4 overexpression drastically increased the epigenetic dysregulations observed upon *Tet2* KO (**Fig.5E**), with upregulated peaks strongly enriched in the stem cell signature and downregulated peaks in myeloid differentiation signatures. This finding was also confirmed by plotting the accessibility of the differential peaks on UMAP coordinates (**Fig.5F**), showing that upregulated peaks have high accessibility in primitive cell types. These data strongly indicate that in the presence of high Sox4 activity, *Tet2* KO cells are likely stalled at primitive stages of differentiation. Finally, this finding was also confirmed by the skewed distribution of Sox4 OE *Tet2* KO cells, which accumulate in the early progenitor stages and are prevented from entering into myeloid commitment as compared to Tet2 KO alone (**Fig.5G**, compare with Fig.1F).

Overall, these data indicate that *Sox4* acts in a synergistic manner with *Tet2*, leading to increased phenotypic and epigenetic dysregulation.

### Cell reprogramming by *Sox4* induces a hypercompetitive HSC state in vivo

To understand the mechanism underlying the observed *Sox4*-mediated increase in stem cell properties and hyperresponsiveness to *Tet2*, we utilized the well established, PVA-based HSC expansion method^71^. Thereby, we could gene modify and analyze HSC ex vivo without extensively compromising their functional properties. Moreover, the inflammation associated with the inducible Mx1 model for Tet2 deletion presented in Figure 5 could be avoided.

Long term HSC (LSK, CD150+ CD48- EPCR+) from *Tet2* fl/fl mice, transduced with Sox4 OE or control and then with Cre-expressing LV to induce Tet2 KO were cultured in PVA (**Fig.6A**). As functional and molecular readouts, cells were analyzed by flow cytometry for HSC markers, harvested for SHARE-seq and transplanted *in vivo*. In accordance with our hypothesis, we were able to observe that the combination of Sox4 OE and *Tet2* KO resulted in a significant accumulation of primitive progenitors (**Fig.6B**). SHARE-seq analysis revealed molecular signatures highly shared with unmanipulated hematopoietic populations^72^, which allowed us to annotate these primitive subsets (**Fig.6C, Fig.S6A**). Gene Set Enrichment Analysis within the HSC cluster demonstrated that among the most deregulated pathways in the presence of Sox4 were Notch, FGF and PI3K (**Fig.6D, Fig.S6B**). These data suggest pleiotropic effects which might promote growth^73–75^. Accordingly, direct comparison of Sox4OE *Tet2* KO cells vs *Tet2* KO cells showed differential expression of *Fgfr2*, *Camk1d* and *Cdk8* genes in HSC and committed compartments (**Fig.S6C**).Corresponding scATAC seq analysis revealed mild changes in accessibility of TF motifs which again pointed to enhanced cell growth of primitive compartment, as marked by increased activity of Atf3^76^ and reduced differentiation capacity at the level of progenitors (as marked by increases in Gata factors and decreased Spib/c) (**Fig.S6D**). To gain more understanding into the enhanced sensitivity to *Tet2* induced by *Sox4*, we performed untargeted metabolomic analyses (**Fig. 6E**). GMP clones were utilized for this analysis since unbiased mass spectrometry methods benefit from higher cell input. We observed that Sox4 overexpression in combination with Tet2 KO increased metabolites associated with purine metabolism, TCA cycle, Phosphoinositide (PI) signaling and glycolysis (**Fig.S6E**). This is consistent with the scRNAseq analysis performed on PVA-expanded HSC, which also indicated an increase in glycolytic gene expression (Fig.6D). Intriguingly, among the most upregulated metabolites with Sox4 overexpression was Hydroxyglutaric acid (2HG) (**Fig.S6F**), an oncometabolite that directly inhibits Tet dioxygenases^77^. Absence of Tet1/Tet3 activity in combination with Tet2 KO was indeed reported to accelerate the leukemogenesis process^78^ and their indirect inhibition by Sox4 may contribute to the highly increased epigenetic dysregulation observed in Sox4 OE *Tet2* KO cells.

**Figure 6.**
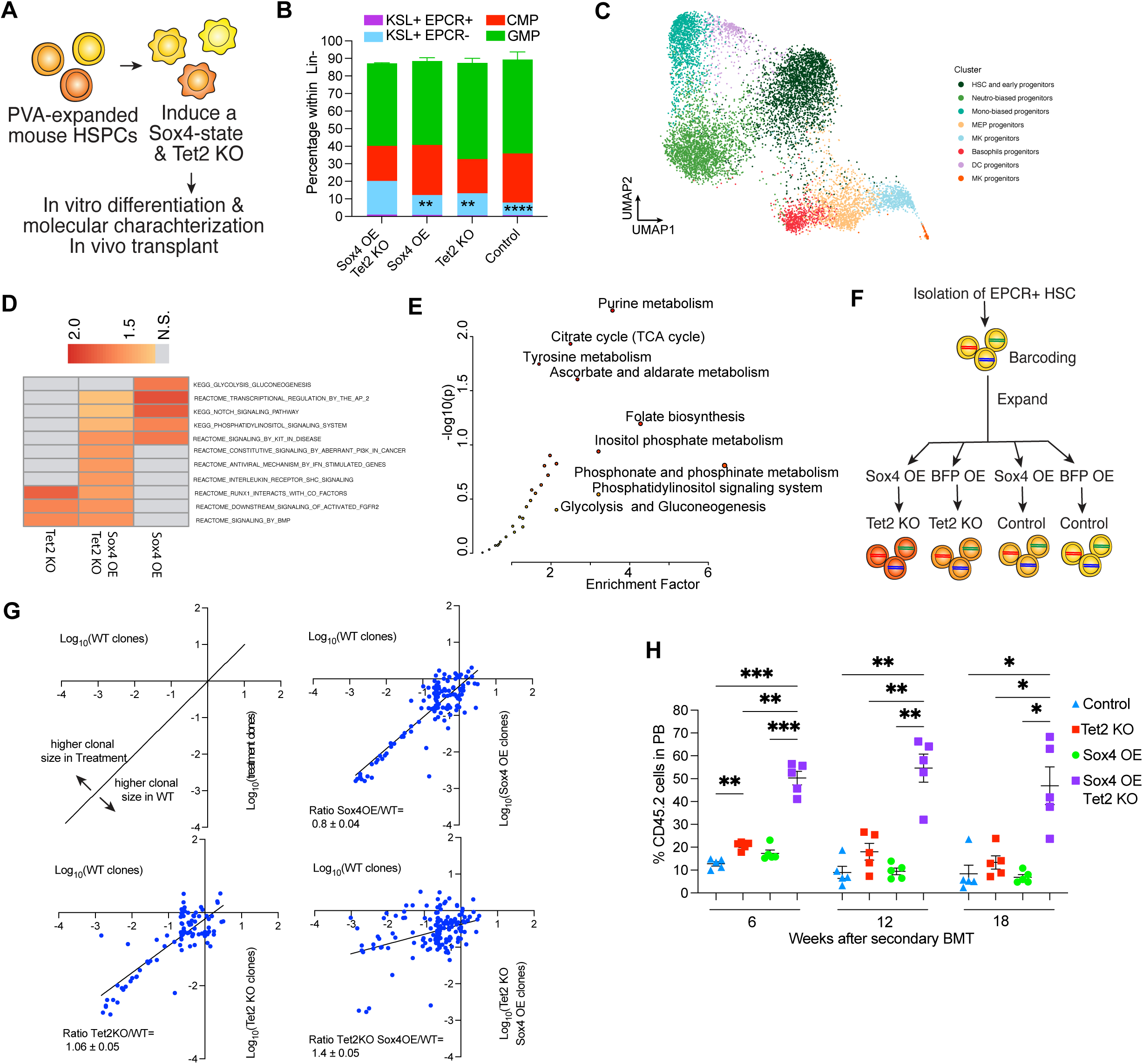
Sox-4 induced cell state provides competitive advantage to *Tet2* KO primitive cells. **A)** Schematics representing the experimental workflow for generating PVA-expanded HSPCs (Sox4 OE and *Tet2* KO) and relative molecular analyses. **B**) Flow cytometry analysis showing the distribution of the indicated primitive population after prolonged expansion of mouse HSC with PVA-containing medium for 3 weeks. n=4, p<0.01, p<0.001. **C**) Unsupervised clustering annotation of ex vivo expanded mouse HSC and analyzed by SHARE-seq **D**) Selected pathways from GSEA analysis comparing gene expression data from HSC cluster from Fig.6C. **E**) Untargeted metabolomic analysis showing deregulated pathways comparing *Tet2* KO Sox4 OE vs WT GMP clones. **F**) Schematics representing the experimental workflow for paired clonal tracking study on PVA-expanded HSPCs. **G**) Clone size comparison of PVA-expanded HSPC sister samples treated as indicated. Clone pairs *Tet2* KO vs WT n=97, Sox4 OE vs WT n= 130, Sox4 OE *Tet2* KO vs WT n= 158. Blue line represents linear regression and highlights relative expansion of Sox4 OE *Tet2* KO clones as compared to WT counterparts, **H**) Percentage of CD45.2 cells measured at different time points in the PB after secondary BM transfer of cells from 6B. n=5, p<0.05, p<0.01, p<0.001.

To test whether Sox4 fosters a hyperproliferative state at a clonal level in primary HSC, we barcoded the cell population using a high-diversity clonal tracing library (**Fig.6F**). After transduction and 7 day PVA expansion the population was split into sister aliquots before introduction of Sox4 OE and *Tet2* KO thereby enabling paired clonal analysis^79^. Comparing the size of paired sister clones across the different treatments, we observed that only the combination of Sox4 OE and *Tet2* KO is able to provide a hyperproliferative advantage compared to WT. Interestingly, only a subset of clones showed this effect further confirming clone-specific behaviors in response to the same genetic modification (**Fig.6G**).

Finally, to validate our findings *in vivo*, we performed competitive transplantation into lethally irradiated mice and compared the hematopoietic output of the genetically modified PVA-expanded HSC. A significant increase in repopulation ability of Sox4OE *Tet2* KO cells was observed on secondary transplantation (**Fig.6H)**. While no changes were noted on primary transplant (**Fig.S6G**), secondary transplant is a reliable assay for HSC self-renewal.

These data demonstrate that *Sox4* activity alters cell sensitivity to *Tet2* mutation in a pleiotropic manner affecting proliferation, differentiation and metabolism to collectively create a cell state of increased competitive advantage.

## Discussion

Somatic mutations in epigenetic regulators have been extensively characterized in mouse models and human samples as drivers of CH. Two questions that are less well defined are approached here.

The first is a fuller definition of the downstream molecular consequences of the mutations which enable clonal outgrowth. It is well known that mutant *TET2* alleles are associated with significant expansion of the myeloid compartments^30^. Here, we defined the multi-omic landscape that underlies differentiation of primitive progenitors towards myeloid lineages and how this network is disrupted in a relevant *Tet2* KO mouse model. In accordance with previous datasets^32,80^, we described that mutant cells display highly overlapping transcriptional and epigenomic profiles as compared to WT controls, highlighting that Tet2-mediated regulation doesn’t globally affect cell identity but rather induces functionally relevant cell states with altered differentiation properties. Consistently, our data indicate that Tet2 regulation is directly required for priming of GMPs towards mature myeloid lineages and in its absence cells accumulate markers of early primitive progenitors such as Runx1, Gata2. This finding might provide a molecular mechanism behind the expansion of GMPs which is observed after *Tet2* KO and provides an enlarged reservoir of progenitors which feeds the myeloproliferation observed especially during inflammatory conditions^46,50^. The functional alterations we found at the level of effector granulocytes or monocytes foster inflammation and decrease pathogen clearance potentially fostering a positive feedback loop.

The second question is how single clones are differentially impacted by a genetically identical mutation. Our data indicate that selected epigenetic states can contribute to clone sensitivity to *Tet2* mutations. By using a clonal GMP system, we were able to correlate the extent of functional changes measured in KO clones to corresponding molecular properties measured in WT counterparts. We demonstrated that chromatin states with features of stem-like cells can prime myeloid progenitors to stronger functional alterations. These data correspond with our finding that *Tet2* regulates stem cell and early progenitor identity before lineage priming, the same setting thought to be the cell of origin in the transition from CH to MDS/AML ^81^. We identified *Sox4* as a driver of hyper-sensitization state towards *Tet2* inactivation. Increased *Sox4* activity is reported to synergize with oncogenes during the process of leukemic transformation^60–63;^ its expression peaks in primitive cells and gradually decreases along myeloid differentiation^35^. Our data are consistent with a model where *Sox4* promotes the maintenance of an immature and hyperproliferative cell state, which increases repopulation advantage and likely the susceptibility of accumulation of secondary mutations. Importantly, this concept was also validated using a clonal tracing method in primary HSC.

We provide evidence that *Sox4* acts through two mechanisms, i) by directly activating transformation-promoting pathways such as Notch, Fgf and PI3k and ii) by potentiating Tet2-mediated epigenetic dysregulation. Increased expansion of *Sox4*, *Hlf* and *Meis1* expressing cells has also been detected in another independent study utilizing inducible *Tet2* KO mice^32^, reinforcing our findings.

Intriguingly, our HSC models highlight that high *Sox4* expression is associated with alterations in genes involved in cell metabolism such as *Cdk8* and untargeted metabolomics findings reinforced this finding. We hypothesize that induction of altered glycolytic states by *Sox4* might play a role in enhancing proliferation/self-renewal of primitive progenitors in CH as described in non-malignant contexts^82^ and during AML development^83^. We thus speculate that metabolic reprogramming may not only be a hallmark of cell transformation but also play an active role in the selection/predisposition of pre-malignant clones for disease development.

Furthermore, our data highlight that individual cell types can be affected by *Tet2* differently. Tet2 interaction with DNA relies on the interaction with chromatin modifiers and TFs, due to the lack of a direct DNA binding domain. This interaction enables highly cell type and chromatin context-specific activity^84^. In hematopoietic cells, CH studies described preferential recruitment of Tet2 at distal enhancer regions in myeloid cells, where it co-localizes with TFs involved in lineage specification and differentiation (Erg/Etv, Runx1-3, Gata1-3, Cebpa/b)^44^. The pleiotropic effects of *Tet2* KO among different cell types could be thus explained by differential availability and DNA binding capacity of its interactors. Methylation in the binding site (and thus reduced DNA accessibility) of key Tet2 interactors has been proposed as a mechanism to explain incorrect lineage differentiation in Tet2 deficient human cells ^85^. Similarly, *Tet2* mutation was reported to synergize with inactivating mutations in key interactors such as PU.1 for leukemic transformation^86^, highlighting the concept that availability of Tet2 interactors might also play a role in determining the downstream functional impact of the mutation.

A potential limitation of our study is the use of an in vitro clonal system for the identification of *Tet2*-related functions. This model may be suboptimal mainly because i) cells are conditionally immortalized using HoxB8, which could influence chromatin accessibility and alter regions bound by Tet2 and ii) cells are arrested at GMP state, therefore missing information regarding more primitive progenitors relevant for disease progression. Complementary *in vivo* studies using clonally tracked HSC-based models such as recently described^27^ could allow uncovering of other molecular players that determine clonal advantage in CH models. Here, we validated the interplay between *Sox4* and *Tet2* in primary HSC-based models and, importantly, other studies indicate that the mechanism could be conserved in humans and relevant for disease progression. Indeed, a recent study describing the molecular landscape of AMLs showed that CTCF binding is perturbed in patients. Aberrant CTCF sites are enriched in SOX4 motifs specifically in *TET2* mutated samples and pathway analysis showed enrichment for Notch and Wnt signaling^87^.

Our study has implications regarding the definition of risk criteria for CH patients. Recent studies exploiting combined single-cell allele genotyping with transcriptomics/epigenomics analyses^80,88,89^ have characterized the landscape of human CH mutant cells. Also, the identification of clonal identities within mutant cells has been accomplished with additional integration of clone-specific molecular identifiers, such as mitochondrial or somatic DNA mutations^90–93^. While these studies highlighted some key molecular differences of mutant populations or clones at different stages of hematopoietic transformation, they have not been able to identify molecular factors that may facilitate the transition from CH to MDS/AML. Here, by using a prospective approach, we uncovered Sox4 expression levels as a specific molecular target in CH mutant cells implicated in increasing clonal selective advantage and expansion. To formally validate this finding in human cells, longitudinal analysis of this biomarker in single CH clones will be required.

The contribution of the epigenome to overt cancer evolution has recently emerged as a highly significant concept^94,95^. For example, a recent study demonstrated that clonally stable and heritable chromatin alterations drive colorectal cancer, and progression is associated with reactivation of homeobox genes^96^. Our work extends for the first time this concept to pre-malignant states and defines that the epigenetic features of the clone in which a mutation occurs can determine the phenotype acquired. For CH, this finding can provide insight into the incomplete penetrance for disease development observed in patients. Further, we identified Sox4 activity as facilitating Tet2-related molecular alterations that foster clonal outgrowth in vivo. Our results validate the hypothesis that epigenetic features can predispose mutant hematopoietic clones for transformation and underscore the importance of pursuing prospective clonal tracking studies for the identification of such risk states for CH malignant evolution.

## Acknowledgments

We thank all the members of Buenrostro and Scadden labs for useful discussion. We also thank David Sykes (MGH) for suggestions related to the Hoxb8 clonal system and Ruslan Soldatov (MSKCC) for critical discussion of the work. We are grateful to the Harvard Stem Cell Institute-Center for Regenerative Medicine Flow Cytometry Core Facility at MGH for technical assistance with FACS sorting, Bauer Core Facility at Harvard University for high-throughput sequencing assistance and Harvard Center for Mass Spectrometry for assistance with metabolomics analyses. G.S. was supported by EMBO long-term fellowship (ALTF-743-2018) and AICF 2018 fellowship. T.A.K. was supported by the Swedish Research Council’s International Postdoc grant. C.M. was supported by a Walter-Benjamin fellowship from the German Research Foundation (Deutsche Forschungsgemeinschaft [DFG], GZ: MA 9452/1-1). C.R. was a recipient of the Kirschstein National Research Service Award (NRSA) Institutional Research Training Grant T32 program.

## Author Contributions

G.S. conceptualized the project, designed and performed experiments, analyzed data and wrote the manuscript. V.K. analyzed data, performed bioinformatic analyses and wrote the manuscript. F.D, T.K. and C.M. designed, performed experiments and analyzed data. R.S, A.E., Y.H. and T.T performed experiments. C.R. helped establish the HOXB8 line. J.B. conceptualized the project, provided supervision, performed bioinformatic analyses and wrote the manuscript. D.T.S conceptualized the project, provided supervision, and wrote the manuscript. All authors read and approved the final manuscript.

---

**Figure S1.**
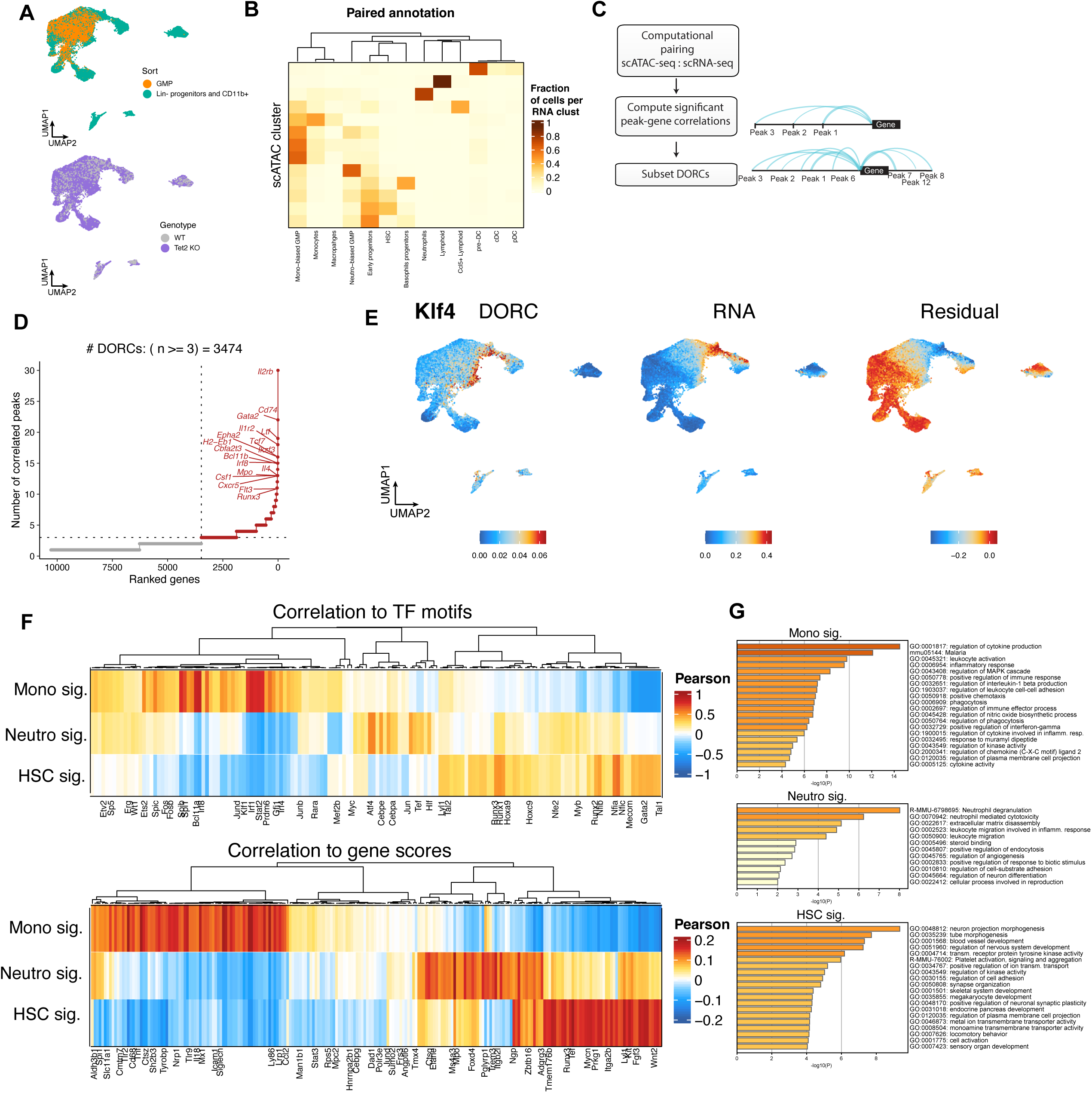
**A**) UMAP plots highlighting distribution of sorted GMP or Lin-/Cd11b populations (top) or genotype (bottom). **B**) Heatmap showing the correlation between unbiased clustering using scATACseq data and unbiased clustering using paired scRNAseq data. scRNAseq data were annotated using key maker gene signatures for each population, as described ^34,35^. Some exemplificative markers include: *Ifitm3*, *Txnip*, *Hlf*, *Adgrl4*, *Procr* (HSC and progenitors), *Vpreb3*, *Cd79a/b*, *FcrIa*, *Ccl5* (lymphoid lineage), *Ly86*, *Klf4*, *Irf8*, *Ccr2*, F13a1, Mpo, C1qb (monocytic/mac lineage), Cebpe, Gstm1, FcnB, S100a8/9, Elane, Lrg1 (granulocytic lineage), *Cd74*, *Itgb7*, *Naaa*, *Cst3*, *Siglech*, *Cd7*, *Ccr7* (DC lineage), *Prss34* (basophilic lineage). **C**) Schematics showing the computational method used for calculation of DORCs **D**) Ranked identity of significant DORCs. **E**) UMAP plots comparing *Klf4* DORC and RNA activity. Plot on the right shows the difference between chromatin accessibility and expression (Residual). **F**) Heatmaps showing Pearson correlation of defined chromatin signatures from Fig.1E to TF motifs (top) and gene scores (bottom). **G**) Enrichment analysis of top correlated gene scores for each chromatin signature.

**Figure S2.**
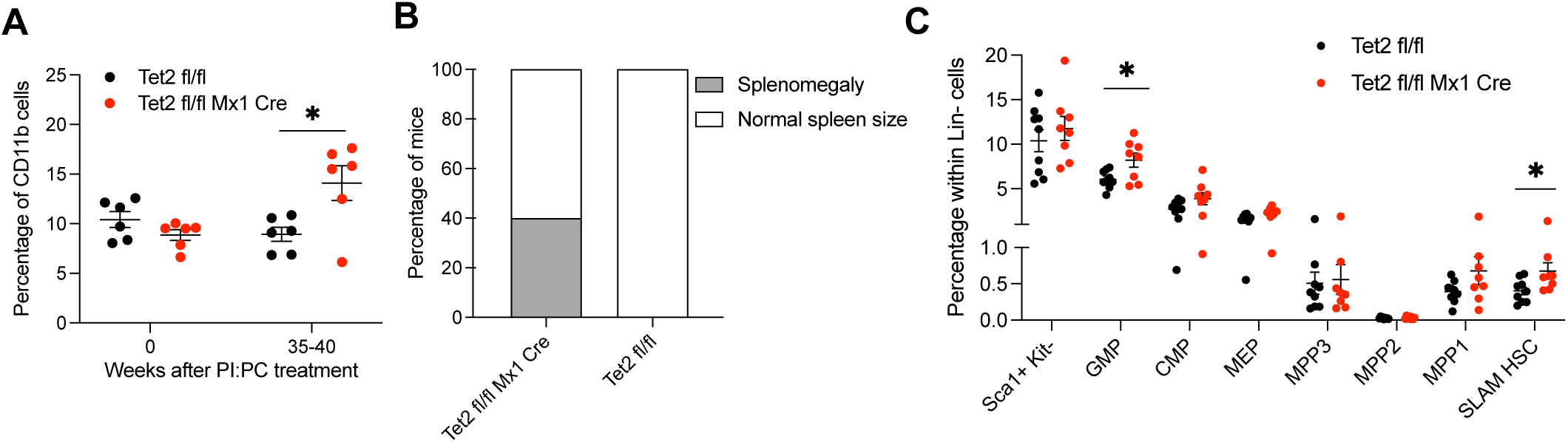
**A**) Percentage of PB CD11b+ myeloid cells before and after *Tet2* deletion by pI:pC treatment. *Tet2* fl/fl n=6; *Tet2* fl/fl Mx1 Cre n=6; p<0.05 **B**) Percentage of mice showing splenic abnormalities stratified by genotype at terminal endpoint of 35-40 weeks after mutation induction *Tet2* fl/fl n=6, *Tet2* fl/fl Mx1 Cre n=6. **C**) percentage of the different BM primitive populations gated within Lin-progenitors.*Tet2* fl/fl n=9; *Tet2* fl/fl Mx1 Cre n=8; p<0.05.

**Figure S3.**
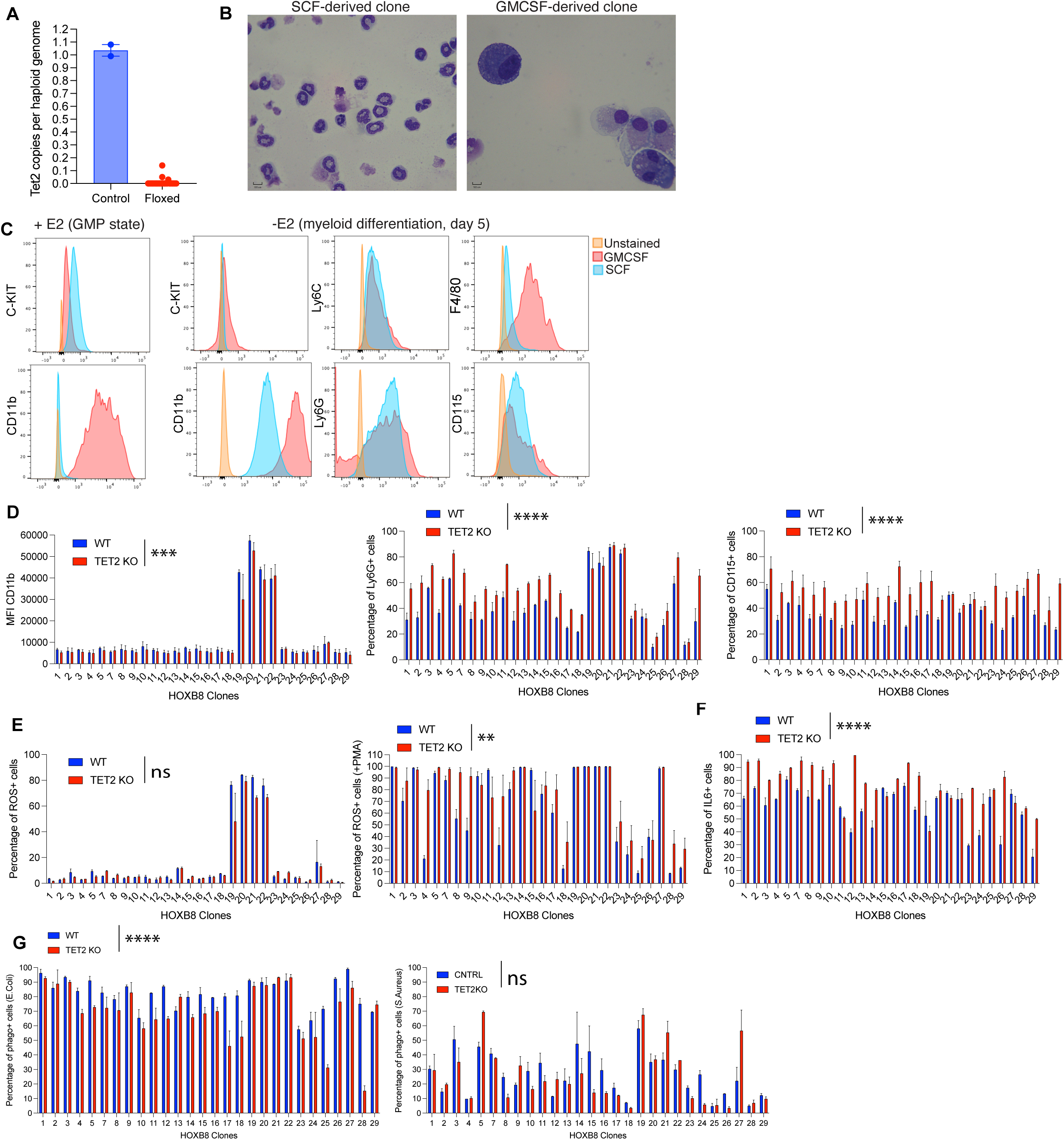
**A**) qPCR showing number of *Tet2* copies each haploid genome in WT and Tet2 KO clones. **B**) Image of cytospin and Wright-Giemsa staining of representative differentiated GMP clones derived using SCF or GMCSF cytokine as indicated at day 5 after estrogen removal. **C**) Representative histograms showing expression of relevant markers on GMP clones derived using SCF or GMCSF cytokine before (+E2) or after differentiation (-E2). Expression levels of Kit indicate GMP state whether absence of Kit and expression of Cd11b indicate more differentiated cell state. **D**) Histograms showing for each matched WT and Tet2 KO GMP clone pair levels of CD11b (left, p<0.001), Ly6G (middle, p<0.0001) and Cd115 (right, p<0.0001) n=3 for each clone. Assay was performed 5 days after clones differentiation. **E**) Histograms showing for each matched WT and *Tet2* KO GMP clone pair levels of ROS expression in a basal state (left) and after stimulation with PMA (right, p<0.01) n=2 for each clone. Assay was performed 5 days after clones differentiation. **F**) Histograms showing for each matched WT and *Tet2* KO GMP clone pair IL6 production after stimulation with LPS (p<0.0001). n=3 for each clone. Assay was performed 5 days after clones differentiation. **G**) Histograms showing for each matched WT and *Tet2* KO GMP clone pair levels of phagocytosis when cells were challenged with E.Coli bioparticles (left, p<0.0001) or with S.Aureus bioparticles (right) after stimulation with PMA (right, p<0.01) n=2 for each clone. Assay was performed 5 days after clones differentiation.

**Figure S4.**
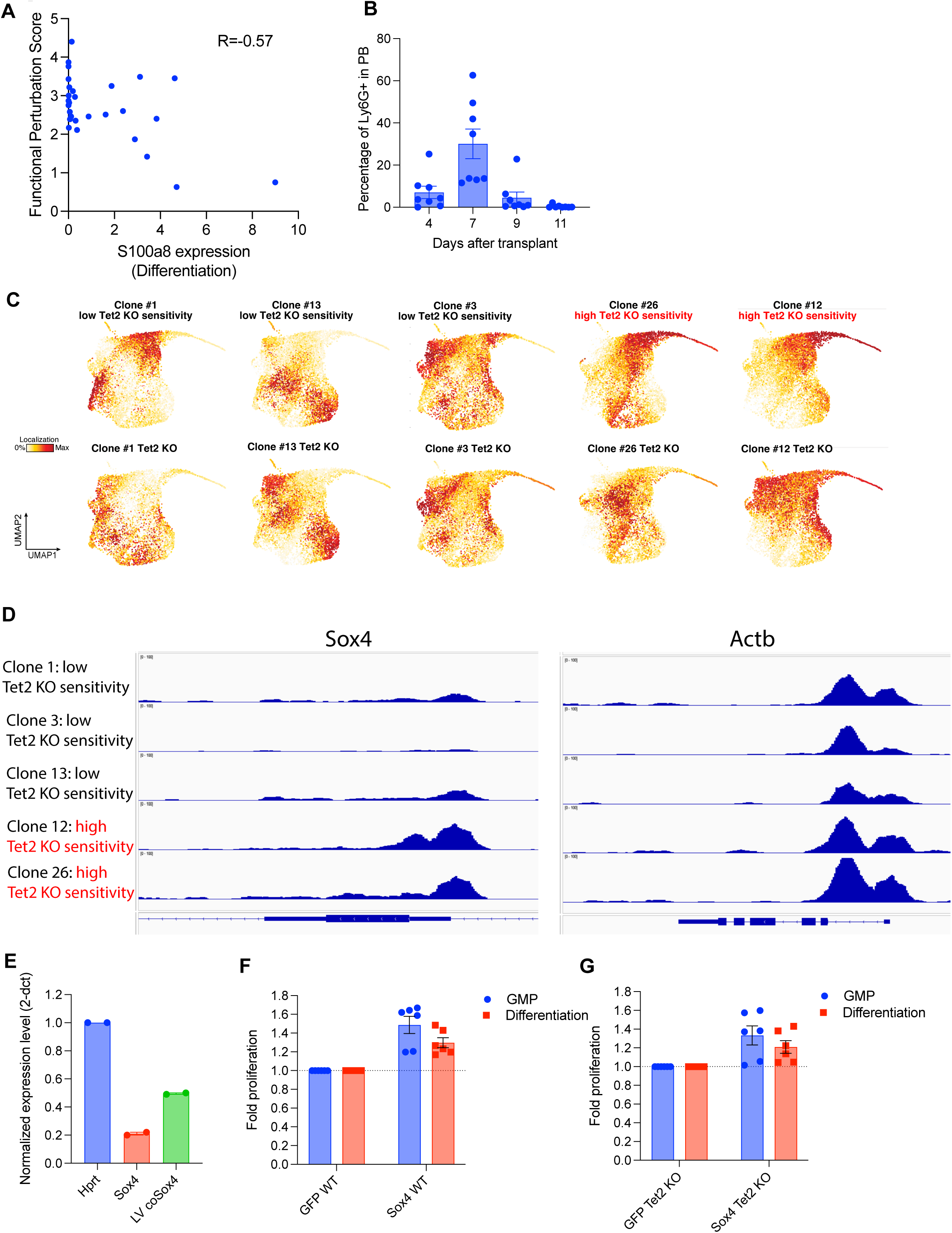
**A**) Pearson Correlation coefficient comparing the Functional Perturbation score and transcript levels of S100a8 in undifferentiated GMP clones. n=28. S100a8 represents a marker of GMP differentiation^53^. **B**) Percent output in the PB at different time points after transplant of SCF GMP clones in lethally irradiated recipient mice. n=8. **C**) UMAP plots showing single cell distribution of WT and mutated GMP clone pairs analyzed by SHARE seq. **D**) Representative ATAC seq tracks showing regulatory region of *Sox4* and *Actb* genes in different GMP clones endowed with different levels of functional alterations observed after Tet2 mutation **E**) Quantitative PCR showing expression of *Sox4* from endogenous transcript (Sox4) or from the overexpressing LV (LV-codon optimized Sox4). Data are normalized to the expression of *Hprt*. **F**) Fold change in the cell number comparing control (GFP overexpressing) and Sox4 overexpressing WT clones.n=6. Measurements are performed 5 days after plating equal cell number under GMP maintaining condition or differentiating conditions. **G**) Same as in B, comparing Tet2 KO clones.n=6.

**Figure S5.**
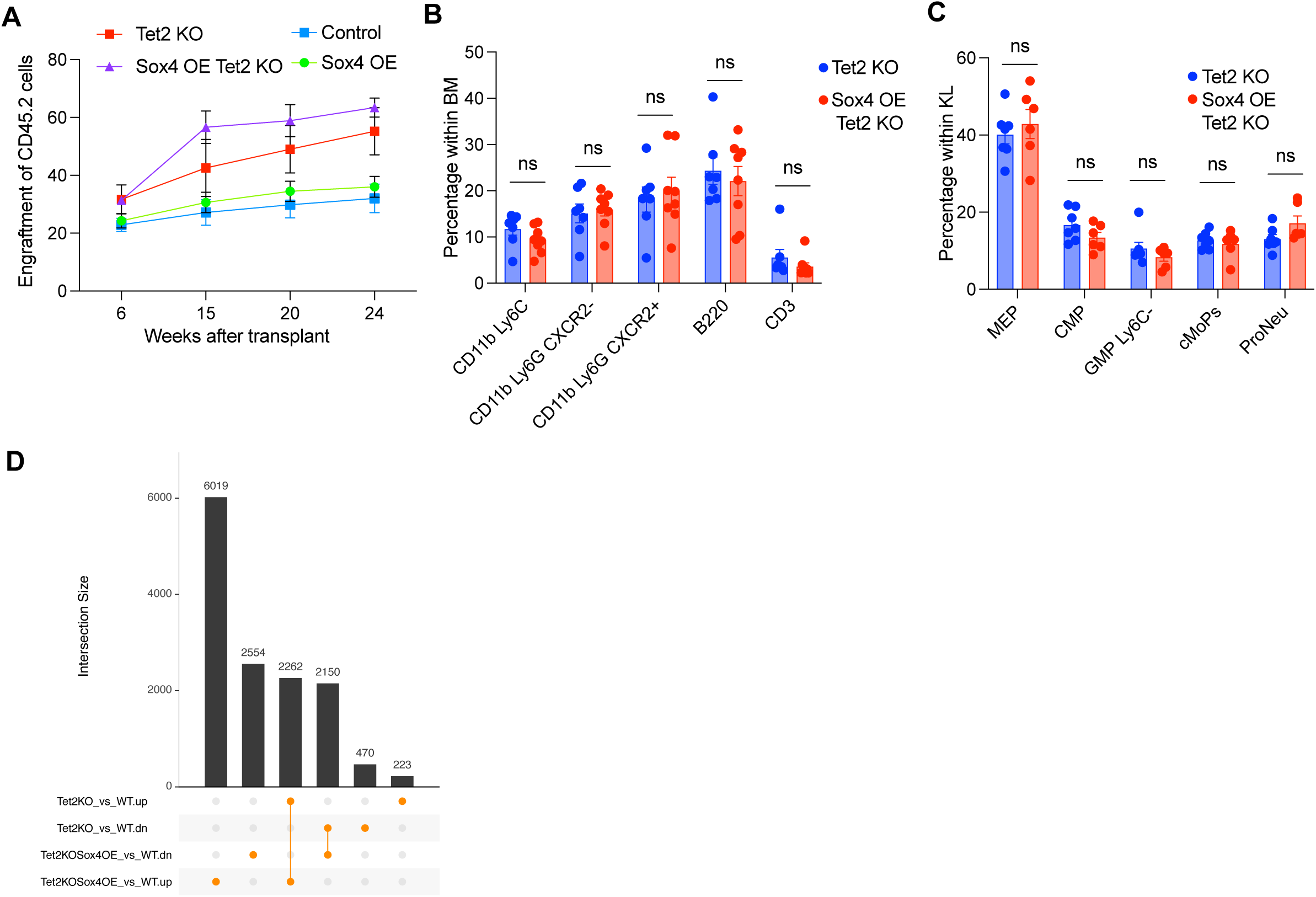
**A**) Percentage of CD45.2 cells measured at different time points in the PB of mice transplanted with GFP overexpression vector in presence (*Tet2* KO) or absence (control) of inducible *Tet2* KO cells as compared to cells with Sox4 overexpression vector in presence (Sox4 OE Tet2 KO) or absence (Sox4 OE) of inducible *Tet2* KO (n=3). To induce *Tet2* deletion, pI:pC was administered 6 weeks after transplant. **B**) Distribution of the indicated differentiated populations in the BM of mice from Fig.6B 35 weeks after transplant (n=7 *Tet2* KO, n=8 Sox4 OE *Tet2* KO). **C**) Distribution of the indicated primitive progenitor populations within Lin-Kit+Sca-cells in the BM of mice from Fig.6B 35 weeks after transplant (n=7 *Tet2* KO, n=8 Sox4 OE *Tet2* KO). Populations are defined as in^41^ **D**) Barplot showing the overlap between differential peaks from scATACseq dataset comparing *Tet2* KO and Sox4 OE *Tet2* KO cells.

**Figure S6.**
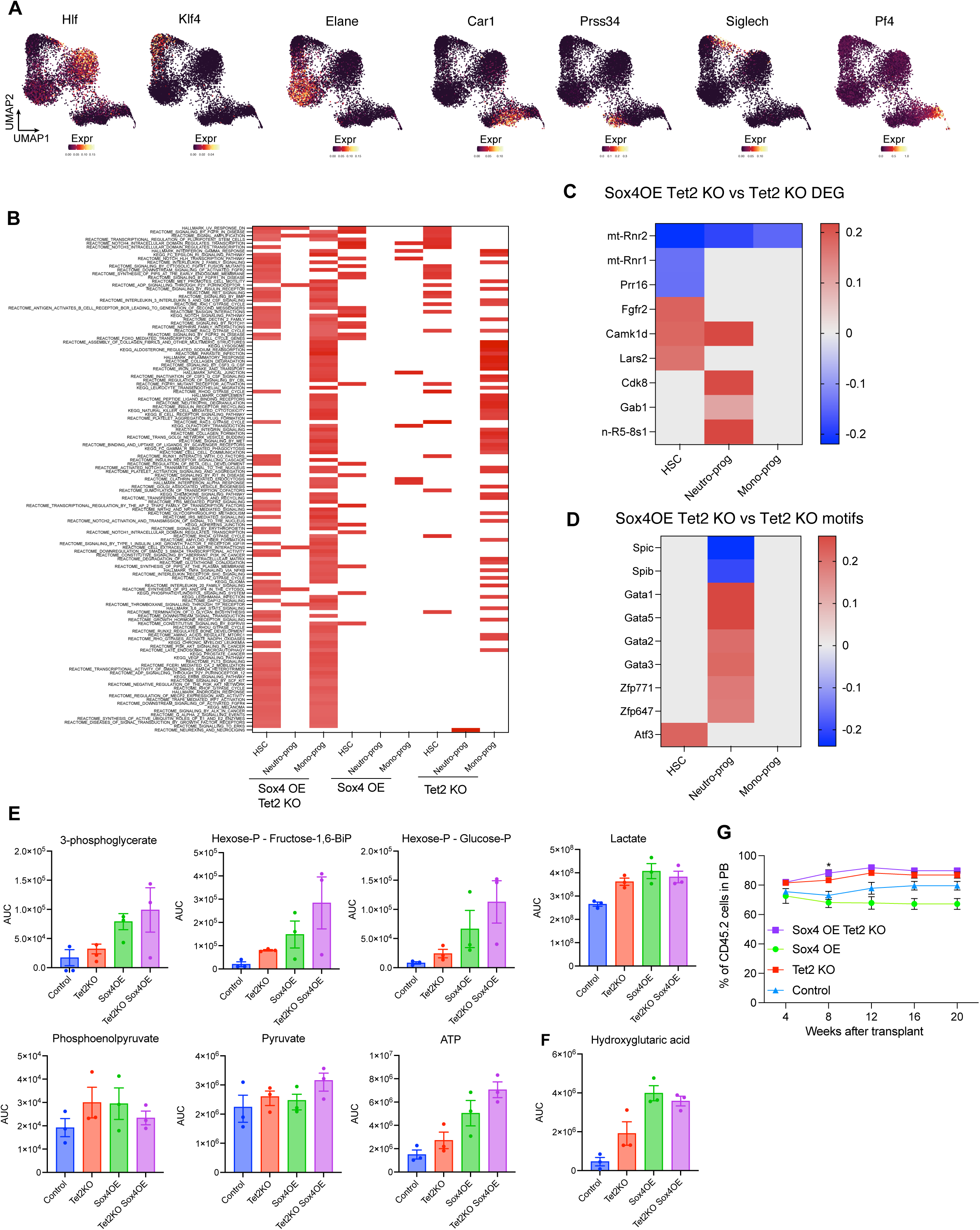
**A**) UMAP plots showing RNA expression of representative markers utilized for annotating PVA-expanded progenitors from Fig.6C. **B**) GSEA analysis comparing gene expression data from Fig.6C. Each sample is compared to WT control cells. HSC cluster, Neutro and Mono progenitor clusters are shown. **C**) Differential genes from Fig.6C (Pval adj<0.05) comparing Sox4 OE *Tet2* KO and *Tet2* KO among the indicated clusters. **D**) TF motif analysis showing Sox4 OE *Tet2* KO vs *Tet2* KO comparison within the indicated cell clusters from Fig.6C. TF motifs with FDR<0.25 are shown. **E**) GMP clones treated as indicated were analyzed by untargeted metabolomics (Fig.6E).n=3. Normalized AUC of different metabolites related to the glycolysis pathway are shown. **F**) Normalized AUC of hydroxyglutaric acid (2-HG) in GMP clones from Fig.6E. **G**) Percentage of CD45.2 cells measured at different time points in the PB after primary BM transfer of cells from 6H .n=3, p<0.05.

## STAR Methods

### RESOURCE AVAILABILITY

#### Lead contact

Further information and requests for resources and reagents should be directed to and will be fulfilled by the lead contact, David T Scadden or Jason D Buenrostro.

#### Materials availability

Plasmids generated in this study are available under a material transfer agreement with Massachusetts General Hospital.

#### Data and code availability

scRNAseq, scATACseq and bulk ATACseq datasets have been deposited at NCBI GEO and will be publicly available as of the date of publication. Accession numbers will be listed in the key resources table. This paper does not report original code. Any additional information required to reanalyze the data reported in this paper is available from the lead contact upon request.

### EXPERIMENTAL MODEL AND SUBJECT DETAILS

#### Mice

*Tet2* fl/fl mice^30^ were obtained from The Jackson Laboratory (#017573) and crossed to Mx1-Cre mice (#003556, The Jackson Laboratory). Mx1-Cre negative littermates were utilized as controls. For bone marrow transplantation experiments, aged-matched CD45.1(STEM) mice^68^ and WT C57BL6/J recipients (#000664, The Jackson Laboratory) were utilized. 8-12 weeks old male and female mice were used in all experiments. All mice were bred and maintained in pathogen-free conditions and all procedures performed were approved by the Institutional Animal Care and Use Committee of Massachusetts General Hospital.

#### Cell lines

HOXB8 immortalized clones were maintained in RPMI medium (Thermo Fisher) supplemented with 1% Penicillin/Streptomycin, 1% Glutamine, 0.5 uM beta-estradiol (Sigma, E2758) and conditioned media containing different cytokines. For the neutrophil-biased cells, media contained approximately 100 ng/ml SCF (generated from a Chinese hamster ovary cell line that stably secretes SCF), whereas for macrophage progenitors GM-CSF 10 ng/ml (Peprotech). HOXB8 cultures were kept in a 5% CO2 humidified atmosphere at 37C.

#### Primary cells

For BM transplant studies, primary mouse Lin-Kit+ Sca+ cells were seeded at the concentration of 5×105 cells/ml in serum-free StemSpan medium (StemCell Technologies) supplemented with 1% Penicillin/Streptomycin, 1% Glutamine, and mouse early-acting cytokines (SCF 100 ng/ml, Flt3-L 100 ng/ml, TPO 50 ng/ml, and IL-6 20 ng/ml; all purchased from Peprotech). For PVA-expansion studies, Lin-Kit+ Sca+ Cd150+ Cd48- Epcr+ cells were seeded in 1X Ham’s F-12 Nutrient Mix liquid media (Gibco), supplemented with 1 M HEPES (Gibco), 1% Penicillin/Streptomycin/Glutamine, 1% ITSX (Gibco), SCF 10 ng/ml, TPO 100 ng/ml (Peprotech), 1 mg/ml PVA (Millipore Sigma). Further details are provided in the following sections. HSC cultures were kept in a 5% CO2 humidified atmosphere at 37C.

## METHOD DETAILS

### scATAC-seq single cell profiling

Cell lysis, tagmentation and droplet library preparation were performed following the SureCell ATAC-Seq Library Prep Kit User Guide (17004620, Bio-Rad). Harvested cells and tagmentation related buffers were chilled on ice. Lysis was performed simultaneously with tagmentation. Washed and pelleted cells were resuspended in Whole Cell Tagmentation Mix containing 0.1% Tween 20, 0.01% Digitonin, 1x PBS supplemented with 0.1% BSA, ATAC Tagmentation Buffer and ATAC Tagmentation Enzyme. Cells were mixed and agitated on a ThermoMixer (Eppendorf) for 30 min at 37C. Tagmented cells were kept on ice prior to encapsulation. Tagmented cells were loaded onto a ddSEQ Single-Cell Isolator (12004336, Bio-Rad). Single-cell ATAC-seq libraries were prepared using the SureCell ATAC-Seq Library Prep Kit (17004620, Bio-Rad) and SureCell ddSEQ Index Kit (12009360, Bio-Rad). Bead barcoding and sample indexing were performed following the standard protocol and the number of amplification cycles was adjusted according to cell input. Libraries were loaded on a NextSeq 550 (Illumina) and sequencing was performed using the NextSeq High Output Kit (150 cycles) and the following read protocol: Read 1 118 cycles, i7 index read 8 cycles, and Read 2 40 cycles. A custom sequencing primer is required for Read 1 (16005986, Bio-Rad).

### scRNA-seq single cell profiling

scRNA-Seq was performed on a Chromium Single-Cell Controller (10X Genomics) using the Chromium Single Cell Reagent Kit v2, Chromium Next GEM Chip A and Chromium i7 Multiplex Kit according to the manufacturer’s instructions. Briefly, single cells were partitioned in Gel Beads in Emulsion (GEMs) and lysed, followed by RNA barcoding, reverse transcription and PCR amplification (according to the available cDNA quantity). scRNA-Seq libraries were prepared according to the manufacturer’s instructions, checked and quantified on Tapestation 4200 (Agilent) and Qubit 4 fluorometer (Invitrogen). Sequencing was performed on a Novaseq machine (Illumina) using the Novaseq S1 Kit (100 cycles).

### Multiplexed Bulk ATAC-seq

Indexed Tn5 transposome complexes were assembled as described previously^38^ Also see this reference for a description of how the barcodes were designed and a table with the oligo sequences. Cells were washed twice with 1x PBS, counted, and resuspended to a concentration of 0.5 x 10^6^ cells/mL. 2 μL of cells in 1x PBS (1,000 cells) were mixed with 2 μL of barcoded Tn5, 5 μL 2x Illumina Tagment DNA Buffer (TD), 0.1 μL 10% NP40 (final concentration 0.1%) and 0.9 μL H2O in a 96 well plate. Each well contained a different sample and Tn5 barcode. Cells were mixed and agitated on a ThermoMixer (Eppendorf) for 30 min at 37C. All the wells were pooled together on ice to prevent cross-contamination between Tn5 barcodes. Tagmented DNA was purified using a MinElute PCR Purification Kit (Qiagen), then minimally amplified for sequencing as previously described^97^. Final libraries were purified using the MinElute PCR Purification Kit (Qiagen), and sequenced on a NextSeq 550 (150 cycles), using the following parameters: Read 1 92 cycles, i7 index read 8 cycles, and Read 2 66 cycles, 50% of PhiX Sequencing Control. A custom sequencing primer is required for Read 1 (16005986, Bio-Rad).

### SHARE-seq

SHARE-seq was performed as described previously^39^. Briefly, single cells were fixed by adding Formaldehyde (28906, ThermoFisher) at a final concentration of 1%. Fixed cells were transposed using barcoded Tn5 (Seqwell) in a transposition buffer (1 × TD buffer from Illumina Nextera kit, 0.1% Tween 20 (P9416, Sigma), 0.01% Digitonin (G9441, Promega)) at 37°C for 30 minutes with shaking at 500 rpm. Transposed cells were reverse transcribed using Maxima H Minus Reverse Transcriptase along with RT primer containing a Unique Molecular Identifier (UMI), a universal ligation overhang and a biotin molecule. Ligation of barcoded adapters was performed using three rounds of split pool barcoding followed by reverse crosslinking. ATAC and RNA libraries were prepared as previously described (Ma et al., 2020). Libraries were quantified with KAPA Library Quantification Kit and pooled for sequencing. Libraries were sequenced on the Nova-seq platform (Illumina) using a 200-cycle S1 kit and the following read protocol: Read 1: 50 cycles, Index 1: 99 cycles, Index 2: 8 cycles, Read 2: 50 cycles.

### scATAC-seq and scRNA-seq data analysis, scATAC-scRNA-seq cell pairing

### scATACseq data processing

Genome-wide chromatin accessibility peaks were called using MACS v2 (MACS2)^98^ on the merged aligned scATAC-seq reads per condition, generating a list of peak summit calls per condition. To generate a non-overlapping set of peaks, we first extended summits of each condition to 800-bp windows (±400 bp). We combined these 800-bp peaks, ranked them by their summit significance value and retained specific non-overlapping peaks on the basis of this ordering. We further added to the peak list all non-overlapping peaks from the ImmGen ATAC-seq atlas, after also extending the ImmGen peaks to 800-bp windows^99^ (https://sharehost.hms.harvard.edu/immgen/ImmGenATAC18_AllOCRsInfo.csv). This resulted in a filtered list of disjoint peaks (n=297,361), which were finally resized to 301 bp (i.e. ±150 bp from each peak summit) and used for all downstream analyses.

### scRNA-seq analysis

Base call files were demultiplexed, for each flow cell directory, into FASTQ files using Cellranger v3.1.0 (https://github.com/10XGenomics/cellranger) mkfastq with default parameters. FASTQ files were then processed using Cellranger count with default parameters. Gene-mapped counts were then loaded into R as a Seurat^100^ object and used for downstream analysis. Genes with at least one UMI across cells were retained, and cells with a number of unique feature counts ≥ 200 and total UMIs ≥ 5000 and mitochondrial read percentage of < 5 % were initially retained. Normalization and scaling of RNA gene expression levels was then performed. PCA dimensionality reduction was run and UMAP was used for the final 2D cell projection (top 30 PCs). A cell kNN graph was determined using the FindNeighbors function in Seurat (*k* = 30 cell neighbors). Cells were then grouped into clusters using the FindClusters Seurat function (resolution = 0.8; Leiden algorithm), and cluster and cell annotations manually assigned by visualizing the mean and percent expression of cell identity markers within cell clusters. Broader annotations were determined by merging finer cell groupings (<u>Fig</u>.S1B).

### scATAC-seq single cell clustering and annotation

First, dimensionality reduction was performed with cisTopic^101^ using the runWarpLDAModels function as part of the cisTopic R package, with the prior number of topics set to 50. Next, Harmony^102^ was run on the cisTopic cell Z-scores to adjust for observed sequencing batch effects (correcting for animal as a batch covariate). The batch-corrected cisTopic cell Z-scores were then used to project cells in 2D by running UMAP as part of the uwot R package, with *k*=50 cell neighbors and a cosine distance metric. Cells were clustered using a Louvain algorithm, and clusters were annotated using gene activity scores and gene expression markers (see below).

### TF motif scores

TF motif accessibility Z-scores were computed for scATAC-seq data using chromVAR^103^. Briefly, scATAC-seq data (*n*=57,232 cells ; n=297,361 peaks) was used as input, and GC bias for each peak was determined using the BSgenome.Mmusculus.UCSC.mm10 reference genome. Mouse cisBP TF motifs (*n*=890 TFs) were then matched against the reference peak set, and *n*=100 background iterations were used, using which deviation Z-scores were estimated using chromVAR’s computeDeviations function.

### Gene TSS activity scores

Gene activity scores based on chromatin accessibility were derived for scATAC-seq data (*n*=57,232 cells) using a sum of accessibility fragment counts around gene transcription start sites (TSSs), weighted inversely to the distance from the TSS, as previously described^43^. Aligned scATAC-seq fragments per cell are weighted based on the inverse distance to gene TSSs, then summed across the chosen window (9,212 bp) reflecting 1% of the total weight for the chosen exponential half-life (1 kb). Gene activity scores were then normalized by dividing by the mean score per cell, and used for downstream analysis.

### Differential peak testing

Differential testing of accessibility peaks was determined using DESeq2^104^. First, only early progenitor cells (excluding terminally differentiated clusters) were retained for differential accessibility signature derivation in progenitor cells. This includes cells annotated as HSC and early progenitors, neutrophil and monocyte-biased GMPs (*n*=22,569 cells). Next, for each annotated cell type, cell peak counts were “pseudobulked” or grouped per mouse sample, annotated cell type and genotype (WT or *Tet2* KO) by summing raw accessibility counts per peak per grouping. DESeq’s negative binomial Wald test was then applied with default parameters, adjusting for celltype as a covariate, yielding estimates of fold-change and significance per peak between Tet2KO and WT samples. Only peaks with adjusted p-value ≤ 0.01 were retained as differentially accessible between the two genotype groups, and were used as signatures for downstream analysis (e.g. TF motif enrichment testing and overlap with Tet2 ChIP-seq peaks). Same analysis was repeated for the Sox4 OE *Tet2* KO group. These peak signatures were also used as peak annotation features with chromVAR to score single cells for KO signature accessibility relative to background peaks (Fig 2B).

### Co-accessibility modules

Chromatin accessibility peak “modules” were derived as previously described^43^. Briefly, TF motif accessibility Z-scores for cells that were annotated as wild-type (WT) GMPs (*n*=8,313 cells) were used. First, TF motifs (*n*=890; see TF motif scores section above) were clustered using a sequence similarity correlation cut-off = 0.8, and then the most variable motif in each cluster was used as a representative TF. Additionally, jackstraw PCA was performed on motif deviation Z-scores to determine TF motifs with significantly variable accessibility using *n*=100 iterations and a jackstraw permutation *P* ≤ 0.05 across the first 10 PCs. This yielded *n*=68 significantly variable TF motif groups. Then, difference in mean accessibility was tested across for all reference peaks (*n*=297,361) using normalized scATAC-seq peak counts (mean-centered per cell) between the TF high vs low cell groups (cell high/low groups divided based on the median Z-score across cells for each TF motif), and significant peaks for any given TF (FDR ≤ 1e-06 two-tailed *t*-test) were retained. Finally, fold-changes of mean accessibility between the high and low groups were used to cluster peaks into co-accessible modules, using a Louvain algorithm for determining communities (*k*=30 peak nearest-neighbors), resulting in distinctly grouped peak modules, which were manually annotated based on the TF motifs that positively or negatively associated with their activity. These were then converted into a reference peak x module binary annotation matrix, and chromVAR was used to compute module accessibility deviation Z scores for each cell in the entire scATAC-seq dataset.

### scATAC-scRNA-seq cell pairing and visualization

Cells were paired between the two modalities using the scOptmatch workflow described previously^105^. Briefly, CCA was first run using Seurat’s RunCCA function ^100^ to co-embed scATAC-seq and scRNA-seq data (using the cell KNN-smoothed normalized scATAC-seq gene TSS activity scores and scRNA-seq gene expression, respectively). Only the union of the top 5,000 variable gene scores (ATAC) and gene expression (RNA) was used to derive the top 30 CCA components, with rescaling of features performed prior to running CCA. These components were then used to pair cells between the two assays based on the minimum geodesic neighbors between ATAC-RNA cells across the entire data. For scATAC-seq cells, gene expression of paired scRNA-seq cells were then used to visualize gene expression markers on the scATAC-seq UMAP.

### Peak-gene cis regulatory correlation analysis

Peak-gene links and domains of regulatory chromatin were determined using the FigR R package^105^. Briefly, using paired scATAC and scRNA-seq data, we determined for each gene a set of *cis*-regulatory peaks that are most correlated with the given gene’s expression. To do this, we tested peaks falling within 10 kb of a given gene TSS for correlation in peak accessibility and paired gene expression across single cells, using *n*=100 permuted background peaks (matched for peak GC-content and mean accessibility) for significance testing. Only peak-gene links with a positive correlation and permutation *P* ≤ 0.05 were retained. DORCs were defined as genes having ≥ 3 significantly associated peaks.

### Pathway enrichment analysis/GSEA

Enrichment analysis was performed using Metascape^106^ using the following ontology sources: GO Biological Processes, GO Molecular Functions and Reactome Gene Sets. All genes in the genome have been used as the enrichment background. Terms with a p-value < 0.01, a minimum count of 3, and an enrichment factor > 1.5 are reported. The most statistically significant term within a cluster is chosen to represent the cluster. For analysis in Fig.S1G, gene scores with correlation >0.1 to each gene module were considered. For analysis in Fig.2F, differential genes with pval<0.001 were considered. For GSEA^107^, we performed pre-ranked analysis (genes ordered by fold-change) using the default settings. MSigDB Hallmark(H), KEGG and Reactome (C2) databases were utilized, pathways FDR<0.25 were considered significant.

### Vectors, plasmids and molecular analyses

Lentiviral constructs expressing GFP, codon-optimized Sox4, or Cre recombinase were cloned into self-inactivating transfer constructs under the expression of human PGK promoter. Lentiviral backbones were obtained through MTA with Naldini lab (SR-TIGET, Milan). Lentiviral vectors were generated using HIV-derived, third-generation plasmids. Stocks were prepared and concentrated as previously described^108^ using HEK-293T(ATCC) as packaging cells. Titration was performed by qPCR using a HEK-293T line with known number of vector integrations per diploid genome as standard. High complexity barcoding library LARRY Barcode Version 1 library^79^ was a gift from Fernando Camargo (Addgene #140024). Retroviral stocks were generated using pCL-Eco plasmid, which was a gift from Inder Verma (Addgene plasmid # 12371) For molecular analyses, genomic DNA was isolated with QIAamp DNA Micro Kit (QIAGEN) according to the number of cells available.

For gene expression analyses, total RNA was extracted using RNeasy Plus Micro Kit (QIAGEN), according to the manufacturer’s instructions and DNase treatment was performed using RNase-free DNase Set (QIAGEN). cDNA was synthesized with SuperScript VILO IV cDNA Synthesis Kit (Invitrogen) and analyzed on a CFX Connect Real-Time PCR System (Biorad). The relative expression of each target gene was first normalized to housekeeping genes and then represented as 2^-DCt for each sample.

### Generation of HOXB8-ER clones

Immortalization of murine BM cells with HoxB8-ER was done as previously described^52^ with the following modifications. Filtered BM cells were layered over Ficoll-Plaque-Plus (GE Healthcare Biosciences), and centrifuged at 400 x g for 25 min at RT without break to enrich for mononuclear cells. Cells were incubated in a 6-well tissue culture plate for 48 hours at 37 ˚C with 5% CO2 prior to the retroviral transduction in RPMI (RPMI-1640, Corning) with 10% 2 mM L-glutamine, and 100U penicillin/streptomycin (all from Thermo Fisher), supplemented with 20 ng/ml stem cell factor (SCF), 10 ng/ml interleukin-3 (IL-3), and 10 ng/ml interleukin-6 (IL-6).Non-adherent cells were harvested and 5e105 cells were plated onto a 12-well tissue culture plate (Corning) coated with 10 mg/ml human fibronectin (Sigma). 1 ml of ecotropic retrovirus encoding MSCVneo-HA-ER-Hoxb8 was applied in the presence of 8 mg/ml polybrene and spinoculation was performed at 1000 x g for 60 min at RT. After transduction cells were maintained in RPMI supplemented with 0.5 uM beta-estradiol (Sigma, E2758) and conditioned media containing approximately 100 ng/ml SCF for the generation of neutrophil-biased cells or GM-CSF 10 ng/ml (Peprotech) for the generation of macrophage progenitors. Antibiotic selection was performed by adding G418 at 1 mg/ml final concentration until untransduced control cells the control were not viable (usually ∼7 days). After selection, cells were FACS sorted into 96 well plates with culture media for the generation of single cell clones.

For generation of *Tet2* KO lines, each clone was transduced with integrase-defective lentiviral vector encoding for Cre recombinase or GFP as control at a Multiplicity of Infection (MOI) of 200.

### Functional assays on myeloid cells

#### Cytospin and Wright-Giemsa staining

Cells were prepared in PBS at a concentration of 1 million cells/ml and spun 1,000 RPM for 60 sec on microscope slides. After air drying for 30 min, slides were sequentially soaked in different dilutions Wright-Giemsa stain (Siemens, 100% 4 min, 20% 12 min, 3X rinse in ddH2O). Coverslips were affixed with Permount Mounting Media (Thermo Fisher) and samples were analyzed at 100X magnification using oil immersion objective.

#### Phagocytosis assay

Cells were pre-stimulated with 100 ng/ml LPS (L2630, Sigma) for 30’, washed in PBS and incubated in Live Imaging Solution (Thermo Fisher) along with labeled E. coli or S. aureus BioParticles (Thermo Fisher, 500µg/ml and 1000µg/ml, respectively) for 1hr at 37°C before flow cytometry analysis.

#### Reactive Oxygen Species (ROS) assay

Cells were incubated using Invitrogen™ CellROX™ Flow Cytometry Assay Kit (Thermo Fisher) at 37°C for 30 min in culture media, with or without 80nM PMA (MIllipore Sigma).

#### Intracellular staining for cytokines

Cells were pre-stimulated with 100 ng/ml LPS together with protein transport inhibitor Golgi Plug (1:1000, BD) for 1hr at 37°C. Surface and intracellular staining was performed using Perm/Fix kit (BD) according to manufacturer’s instructions.

#### Proliferation assay

Cells were stained with 0.5µM CellTrace Far Red Cell Proliferation Kit (Thermo Fisher) according to manufacturer’s instructions. Flow cytometry analysis was performed 3 days later.

#### In vivo transplant

20 million HOXB8-ER GMPs were transplanted in lethally irradiated recipients (9.5Gy) together with 200K supporter CD45-mismatched BM cells via retro-orbital injection. Peripheral blood was collected at 4-7-9-11 days after injection.

#### Functional perturbation score

Raw outputs from the functional data were first normalized and the euclidean distance between each WT-KO clone pair was calculated using Factoextra R package.

### Mice

*Tet2* fl/fl mice^30^ (#017573, The Jackson Laboratory) were crossed to Mx1-Cre mice (#003556, The Jackson Laboratory). Mx1-Cre negative littermates were utilized as controls. Genotyping was performed with primers listed in Table 1 (WT amplicon: 250bp, fl/fl amplicon: ∼450bp). For competitive bone marrow transplantation experiments, aged-matched CD45.1(STEM) mice^68^ were utilized. WT C57BL6/J recipients (#000664, The Jackson Laboratory) were subjected to whole body irradiation (9.5Gy) from a 137Cs source one day before BM transplantation. 4 doses of 12.5 ug/g pI:pC (Amersham) were administered intraperitoneally to induce Cre activity.

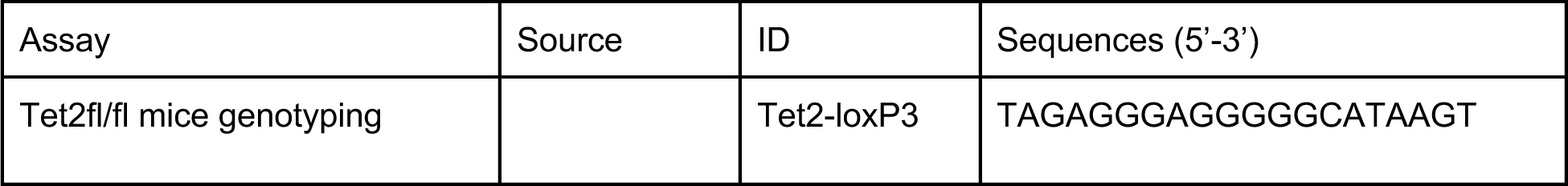
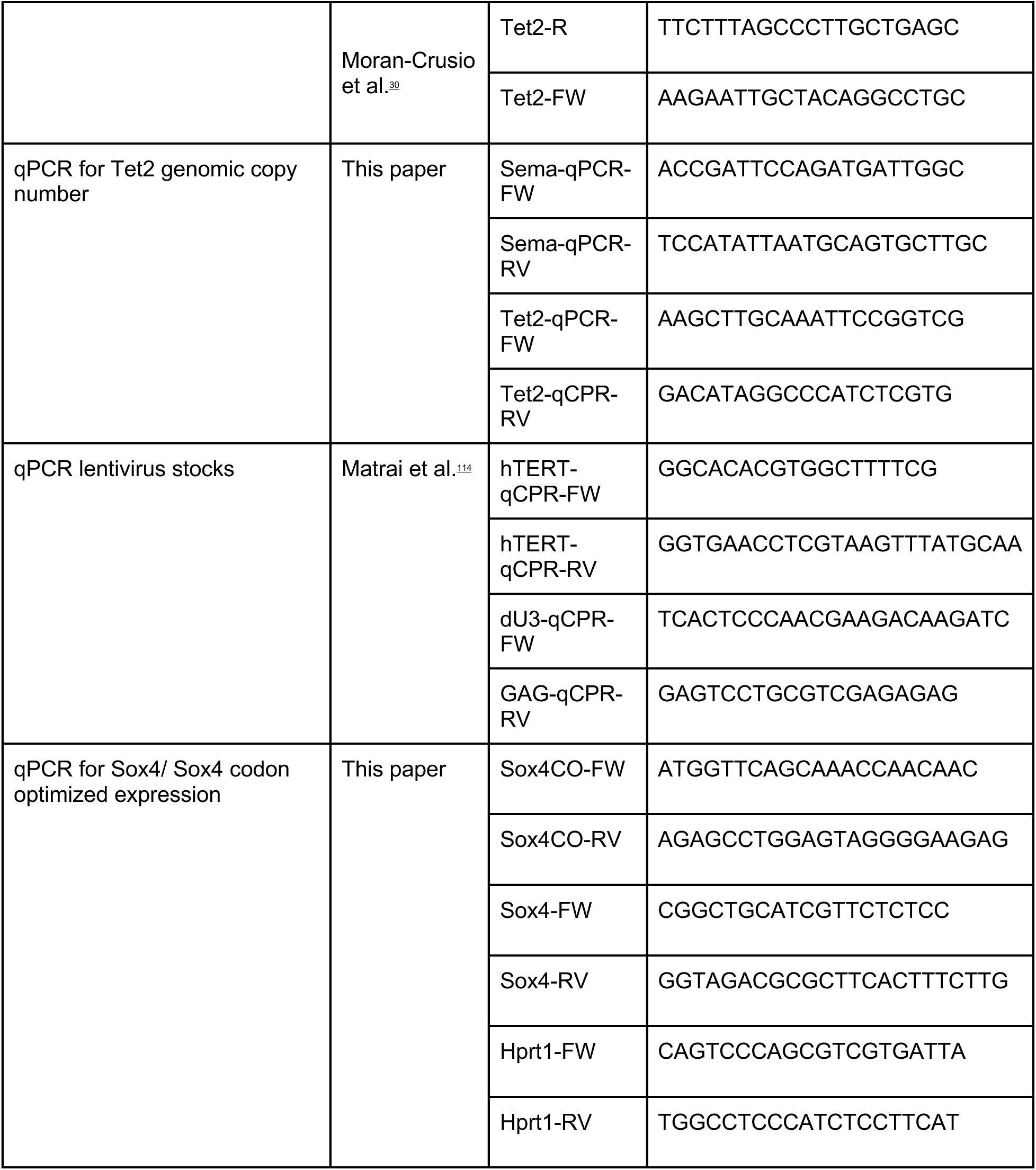
Table 1.

### Bone marrow transplantation of KSL HSPC

Long bones, pelvis and spines were harvested and muscle tissue was removed. Bones were crushed in PBS complemented with 2mM EDTA (Sigma) and 0.5% BSA (Sigma) and bone marrow cells in suspension were filtered on a 40 um cell strainer. Lin-cells were obtained using the Direct Lineage Cell Depletion Kit (Miltenyi Biotec) according to the manufacturer’s instructions. Cells were then stained with antibodies against HSC-related markers Kit and Lin-Kit+ Sca+ (KSL) cells were sorted from *Tet2* fl/fl Mx-1 Cre mice.

Transduction of mouse HSC was performed in serum free-medium enriched with cytokines as previously described^109^ at a MOI of 20. Transduction efficiency was monitored by qPCR. 16 hours after transduction cells were washed in PBS and transplanted at a dose of 10K cells via retro-orbital injection in lethally irradiated C57B6 recipients together with 200K Sca-depleted supporter BM cells. 6 weeks after transplant, 4 doses of 12.5 ug/g pI:pC (Amersham) were administered intraperitoneally to induce Cre activity (and thus Tet2 deletion).

Mice were monitored weekly for body weight and signs of suffering, and euthanized when showing ≥15% weight loss and/or labored breathing, followed by necropsy analysis. Serial collections of blood from the mouse tail were performed to monitor the hematological parameters and donor cell engraftment. At the end of the experiment (25 weeks), BM and spleen were harvested and analyzed (scATAC-seq, flow cytometry for hematopoietic subpopulations).

### PVA-based HSC cultures and analyses

For PVA-expansion experiments, Lin-Kit+ Sca+ Cd150+ Cd48- Epcr+ cells were sorted from *Tet2* fl/fl mice. Culture was performed as previously described^71^. After 6 days of expansion, cultures were split, and transduced with Sox4 OE lentiviral vector or BFP control vector (MOI 40). 4 days later, cultures were split again and transduced with Cre-expressing lentiviral vector (MOI 40) to introduce Tet2 KO. Levels of transduction were monitored by BFP expression and qPCR. LSK cells were enriched again by FACS before proceeding with other analyses (SHARE-seq, flow cytometry for HSC markers, transplant). For transplant, 60K cells from each condition were transplanted in lethally irradiated C57B6 recipients together with 200K competitor total BM cells (mismatched for CD45 isoform expression). Serial collections of blood from the mouse tail were performed to monitor donor cell engraftment. At 30 weeks after primary transplant, 2M cells from each primary mouse were transplanted in secondary lethally irradiated recipients.

For experiments including the high complexity barcoding library LARRY^79^, transduction of Lin-Kit+ Sca+ Cd150+ Cd48- Epcr+ cells was performed 2 hours after sorting in a serum free-medium enriched with cytokines^109^ at a MOI of 20. Transduction level around ∼20% was achieved using these conditions, thus maximizing the likelihood of vector copy number of 1 (1 unique barcode/ cell). 16 hours after transduction, cells were washed in PBS and switched to PVA-based medium. Larry lentiviral libraries were prepared from plasmid stocks as described^79^, and diversity was confirmed to be in the range of 2e5. Libraries were prepared as described^79^, and sequenced on 2×150 Miseq (Illumina). Clonal abundances were estimated using a pipeline adapted from ^110^. Briefly, barcodes are isolated by the identification of flanking sequences using the ShortReads R package, and further filtered by perfect matching of the constant bases present within the 28-mer barcode. Correction for sequencing errors is performed using the Starcode algorithm^111^ using default parameters. Low-frequency barcodes with counts <10 are removed.

### Flow cytometry and FACS

Immunophenotypic analyses and cell sorting were performed on FACS Aria II (BD Biosciences) and antibodies utilized are listed in Key Resource Table. Single stained and Fluorescence Minus One stained cells were used as controls. For peripheral blood analysis, red blood cells were lysed using ACK buffer (Quality Biologicals) for 7 minutes at room temperature before staining. Samples were incubated with the antibody cocktail in PBS 2% FBS for 30 minutes at 4 degrees before analysis or sorting. 7-AAD Viability Staining Solution (BioLegend) was included in the sample preparation for flow cytometry to exclude dead cells from the analysis.

### Metabolomics Analysis

GMP clones were lysed in 100 µl ice-cold 80% methanol in water and polar metabolites were extracted using methanol-chloroform phase separation (1 ml methanol containing 2.5 µM of an internal standard (13C-, 15N-labeled amino acid mix; Cambridge Isotope Laboratories), 500 µl water, and 1 ml chloroform). Samples were then dried under nitrogen flow, resuspended in 20-100 µl 70% acetonitrile in water and run on a ThermoFisher Q-Exactive Orbitrap mass spectrometer equipped with Zic-pHILIC column (150×2.1 mm, 5 mm; Merck). A volume of 5 µl was injected and metabolites were monitored in full-scan, polarity-switching, mode (0 to 45 min, resolution 120,000, RF lens 30%, normalized AGC target 25%, max IT 50ms, m/z range 65 to 1000). Mobile phase A for chromatography consisted of 20 mM. ammonium carbonate, 0.1% ammonium hydroxide, in water and mobile phase B of 97% acetonitrile in water. For untargeted metabolomics, a pooled sample was created from all samples and used for MS/MS runs. For targeted analysis, a standard mix at 1 µM of each compound of interest was prepared and run after the samples to confirm retention times. Metabolite measurements were normalized to the internal 13C/15N-labelled amino acid standard. For untargeted metabolomics functional analysis, peak intensity data (numeric mass (m/z) values) were processed using Metaboanalyst 5.0 software^112^. Positive ion mode, 5.0 ppm tolerance and retention time (minutes) were provided as input parameters. Mouse KEGG database was used for Mummichog Pathway Analysis.

### QUANTIFICATION AND STATISTICAL ANALYSIS

Data were expressed as means ± SEM or dot plots with median values indicated as a line. Statistical tests and number of replicates are reported in the figure legends. Assumptions for the correct application of standard parametric procedures were checked (e.g., normality of the data). Adjusted *p-value*s using Bonferroni’s correction are reported. Whenever these assumptions were not met, nonparametric statistical tests were performed. In particular, Mann-Whitney test was performed to compare two independent groups. In presence of more than two independent groups, Kruskal-Wallis test was performed, followed by post hoc pairwise comparisons. For paired observations, Wilcoxon matched-pairs signed rank test was performed. Analyses were performed using GraphPad Prism 9 and R statistical software. Differences were considered statistically significant at ∗p < 0.05, ∗∗p < 0.01, ∗∗∗p < 0.001, ∗∗∗∗p < 0.0001, ‘‘ns’’ represents non significance.

### KEY RESOURCES TABLE

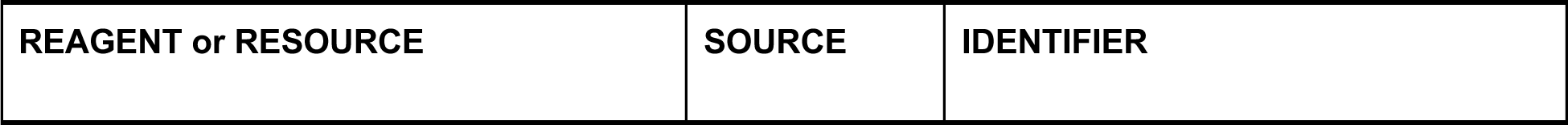
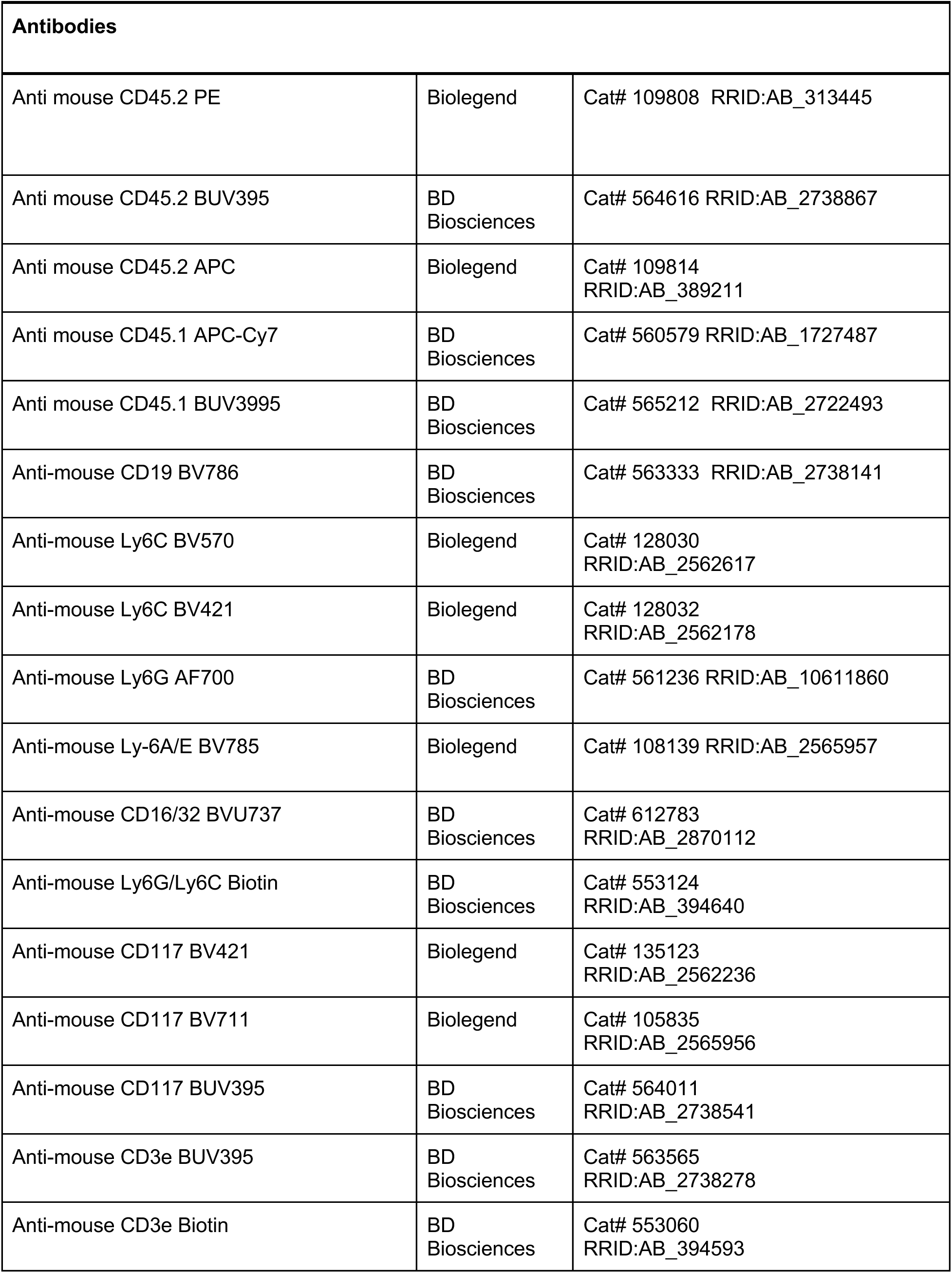
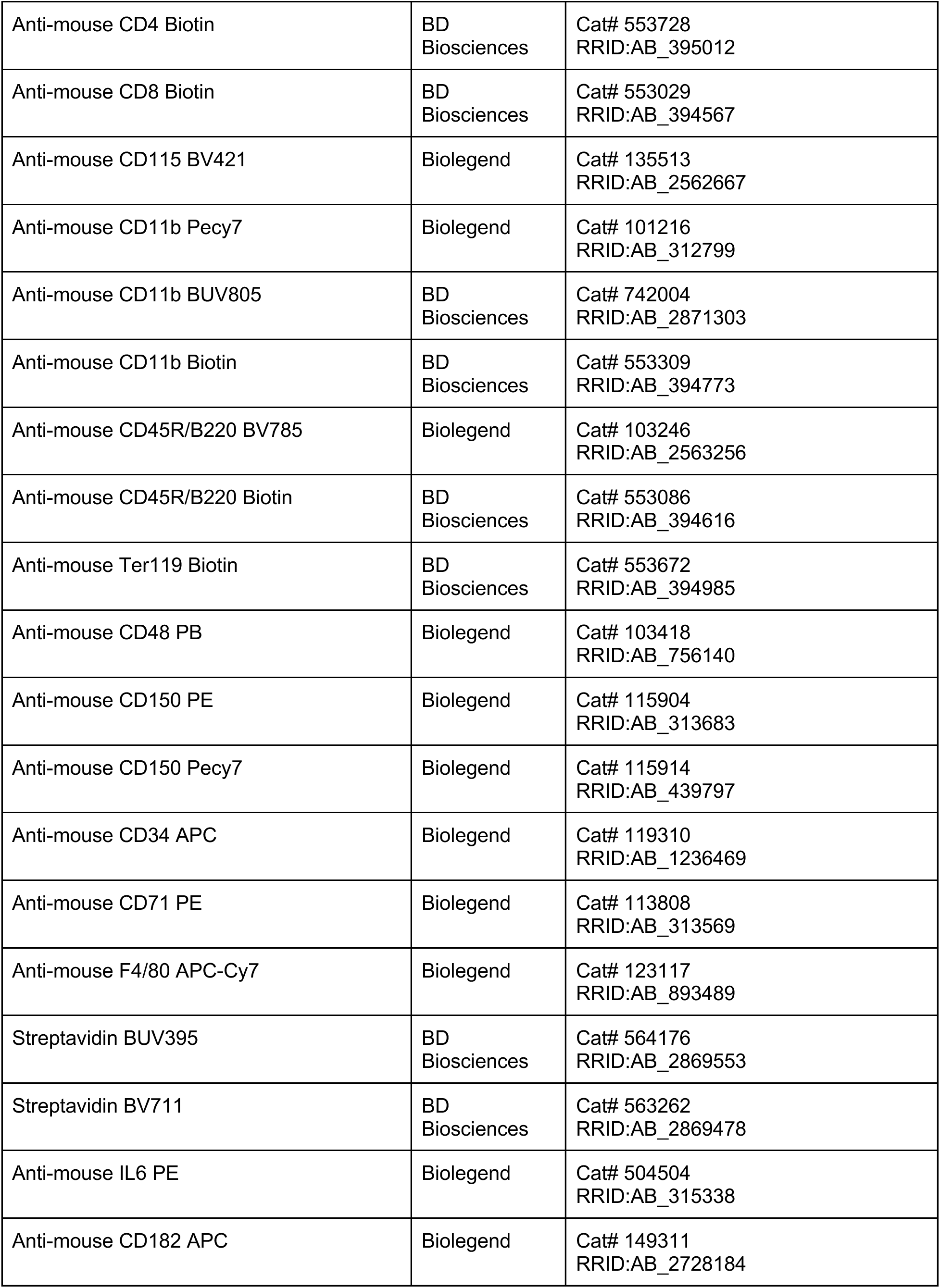
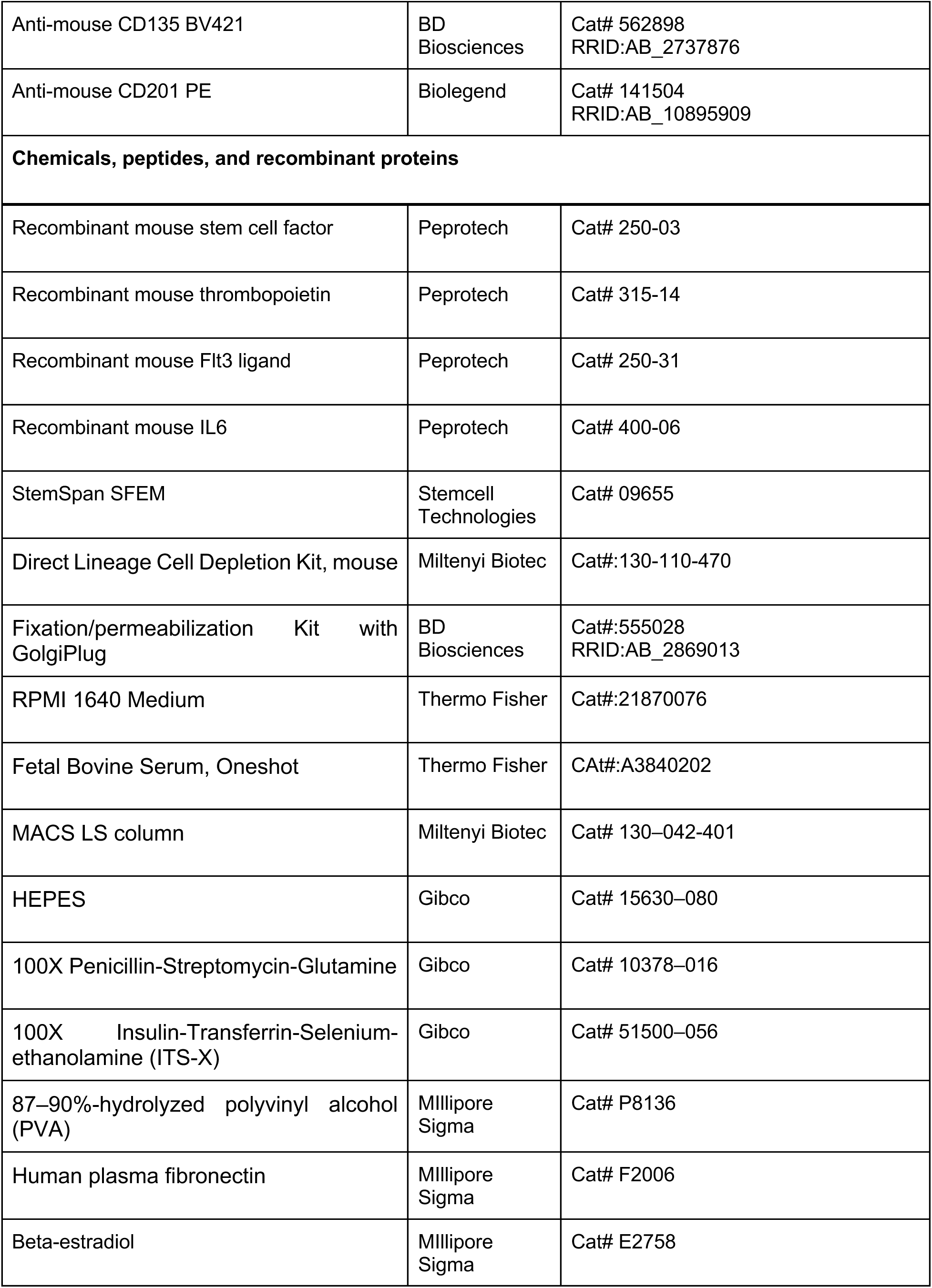
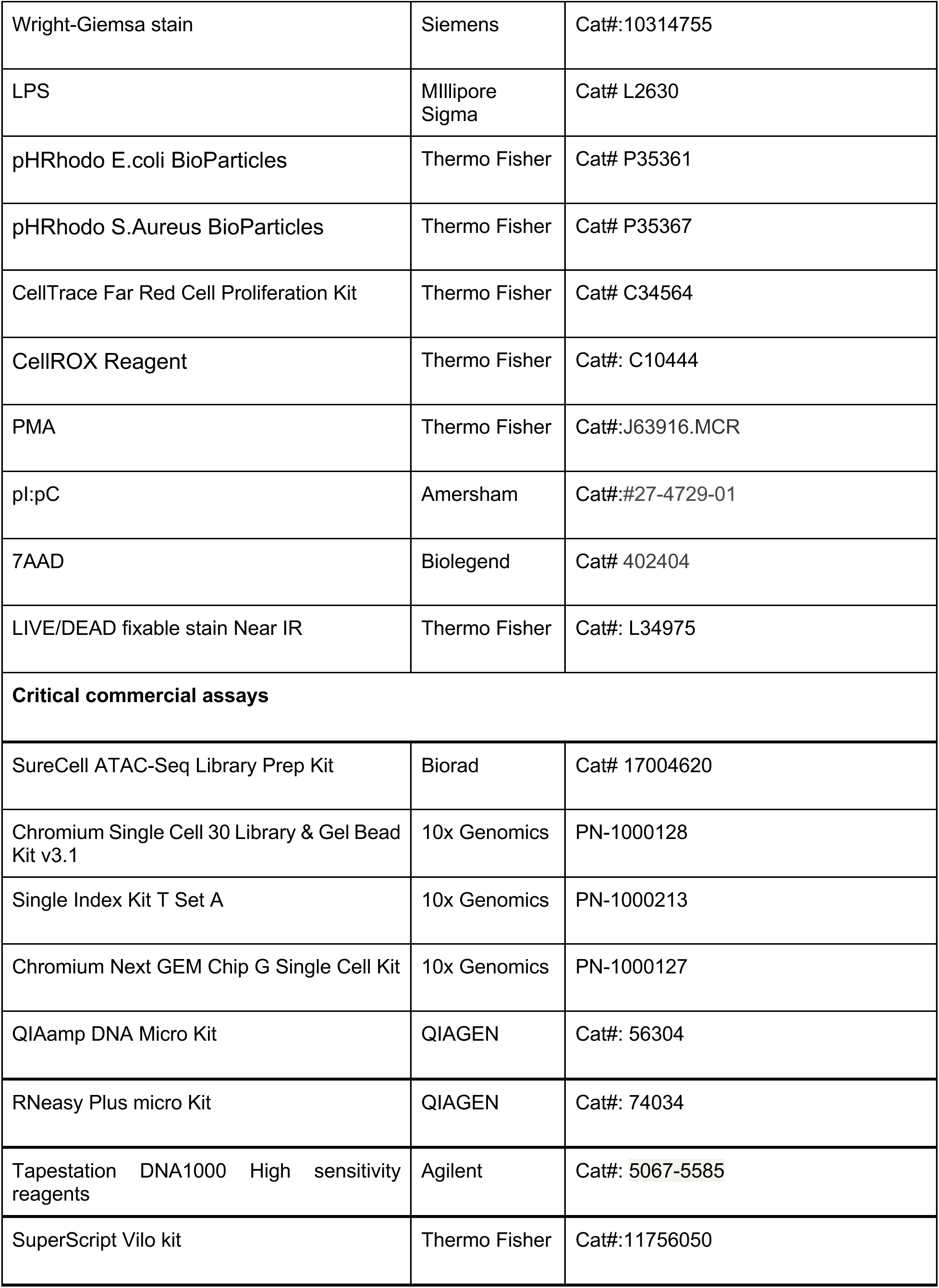
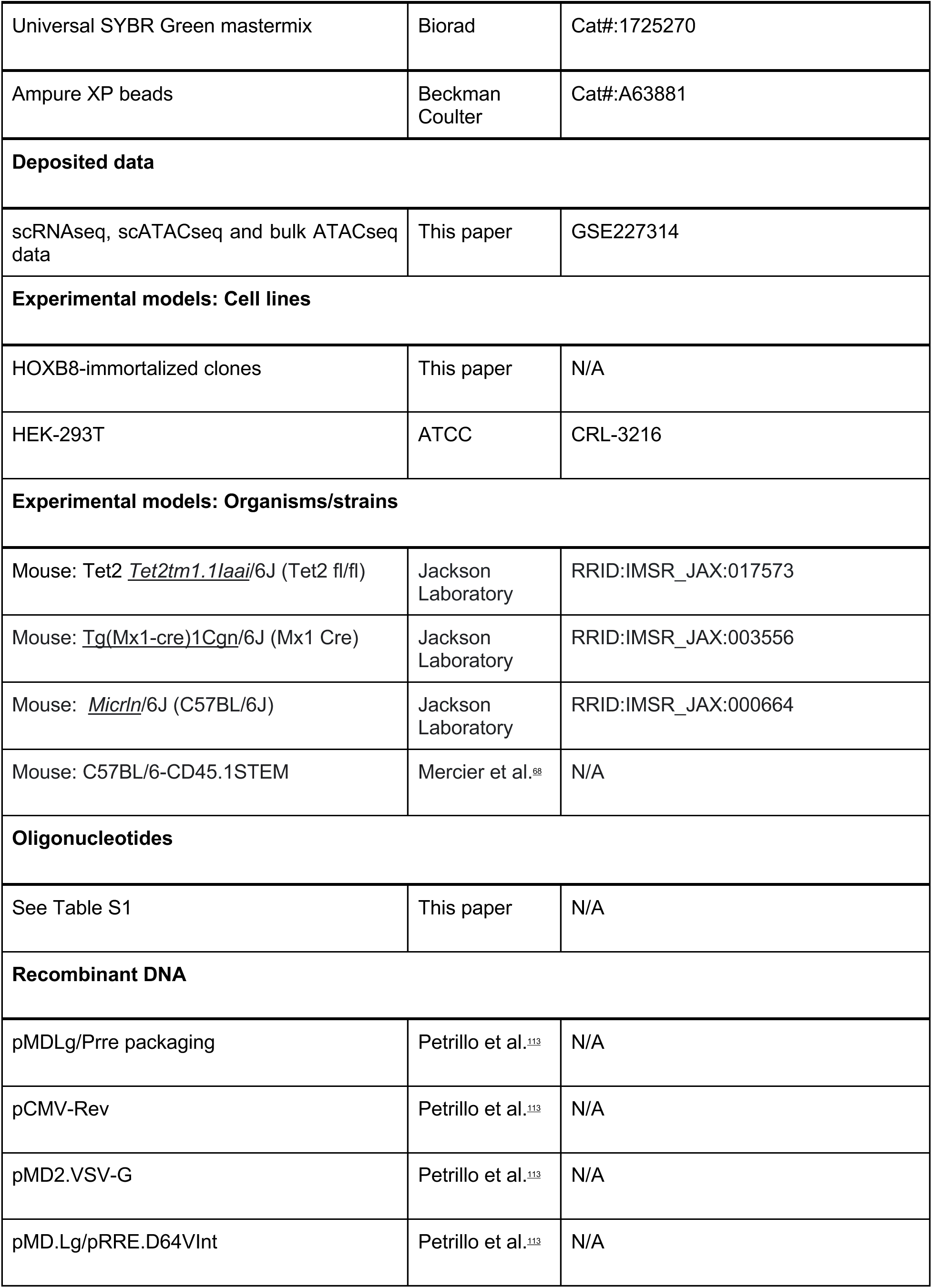
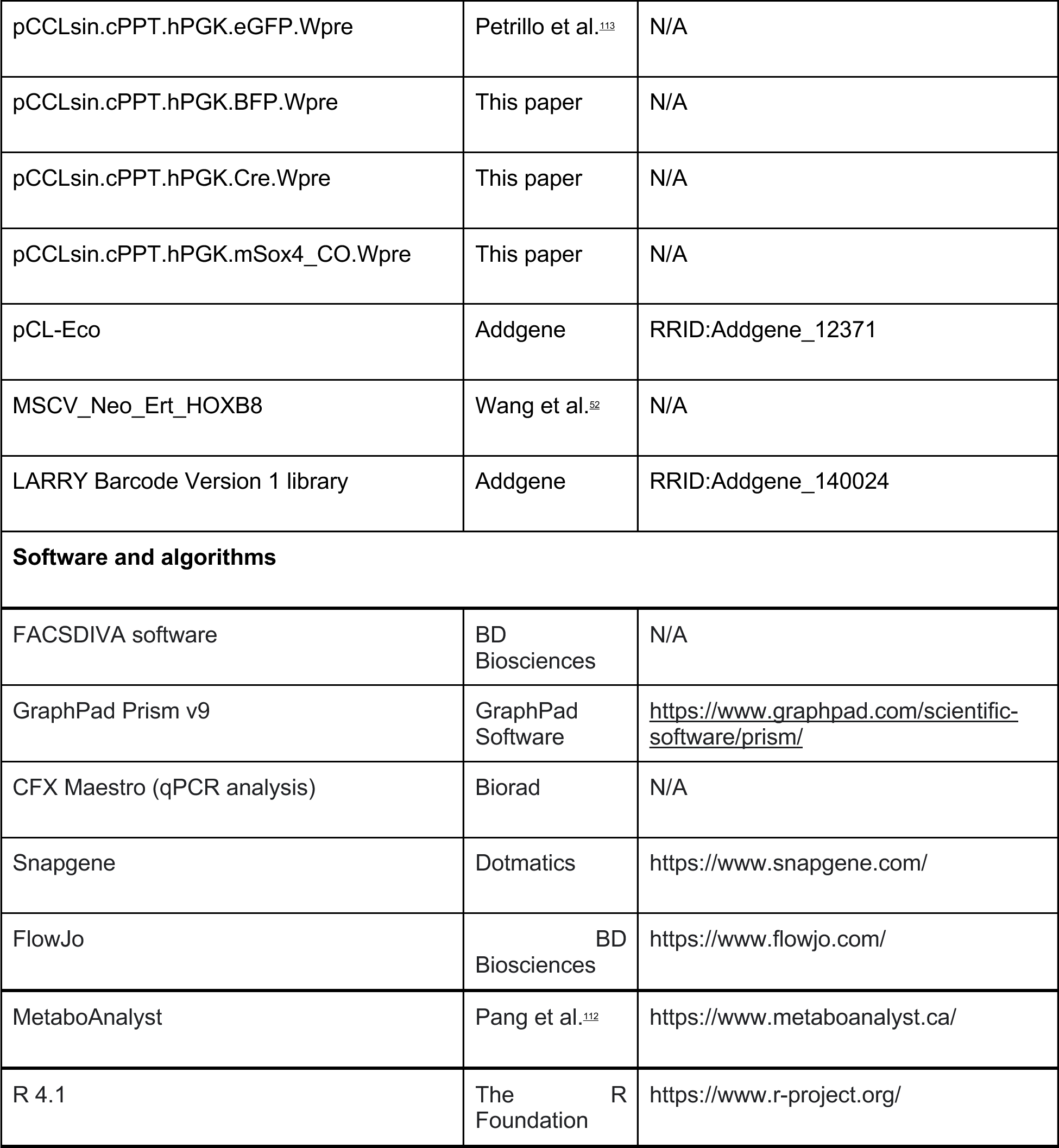

